# Pericentromeric heterochromatin is hierarchically organized and spatially contacts H3K9me2 islands in euchromatin

**DOI:** 10.1101/525873

**Authors:** Yuh Chwen G. Lee, Yuki Ogiyama, Nuno M. C. Martins, Brian J. Beliveau, David Acevedo, C.-ting Wu, Giacomo Cavalli, Gary H. Karpen

## Abstract

Membraneless pericentromeric heterochromatin (PCH) domains play vital roles in chromosome dynamics and genome stability. However, our current understanding of 3D genome organization does not include PCH domains because of technical challenges associated with repetitive sequences enriched in PCH genomic regions. We investigated the 3D architecture of *Drosophila melanogaster* PCH domains and their spatial associations with euchromatic genome by developing a novel analysis method that incorporates genome-wide Hi-C reads originating from PCH DNA. Combined with cytogenetic analysis, we reveal a hierarchical organization of the PCH domains into distinct “territories.” Strikingly, H3K9me2/3-enriched regions embedded in the euchromatic genome show prevalent 3D interactions with the PCH domain. These spatial contacts require H3K9me2/3 enrichment, are likely mediated by liquid-liquid phase separation, and may influence organismal fitness. Our findings have important implications for how PCH architecture influences the function and evolution of both repetitive heterochromatin and the gene-rich euchromatin.

**Author summary:** The three dimensional (3D) organization of genomes in cell nuclei can influence a wide variety of genome functions. However, most of our understanding of this critical architecture has been limited to the gene-rich euchromatin, and largely ignores the gene-poor and repeat-rich pericentromeric heterochromatin, or PCH. PCH comprises large part of most eukaryotic genomes, forms 3D PCH domains in nuclei, and plays vital role in chromosome dynamics and genome stability. In this study, we developed a new method that overcomes the technical challenges imposed by the highly repetitive PCH DNA, and generated a comprehensive picture of its 3D organization. Combined with image analyses, we revealed a hierarchical organization of the PCH domains. Surprisingly, we showed that distant euchromatic regions enriched for repressive epigenetic marks also dynamically interact with the main PCH domains. These 3D interactions are mediated by liquid-liquid phase separation mechanisms, similar to how oil and vinegar separate in salad dressing, and can influence the fitness of individuals. Our discoveries have strong implications for how seemingly “junk” DNA could impact functions in the gene-rich euchromatin.

## Introduction

Nuclear architecture and dynamics regulate many important genome functions (reviewed in [1–4]). The development of Hi-C, which combines chromosome conformation capture (3C) [5] with genome-wide sequencing [6], has led to major breakthroughs in our understanding of global nuclear architecture (reviewed in [7]). However, analyses of Hi-C results have focused on single copy sequences in euchromatic regions (e.g. [6,8–10]), and virtually all have excluded the large Peri-Centromeric Heterochromatin (PCH) portion of genomes due to its enrichment for large blocks of repetitive DNAs [11, 12]. Despite being gene-poor, the PCH plays vital roles in chromosome dynamics [13, 14] and genome integrity [15–17].

A defining characteristic of heterochromatin is its enrichment for ‘repressive’ epigenetic features, such as Histone H3 lysine 9 di- and trimethylation (H3K9me2/3) and its reader protein, Heterochromatin Protein 1a (HP1a) [18, 19]. Interestingly, PCH DNA/chromatin from different chromosomes coalesce into one or a few membraneless PCH ‘domains’ (or chromocenters) in the 3D cell nucleus [20, 21]. Recent studies have shown that specific biophysical properties of HP1a and liquid-liquid phase separation (LLPS) may mediate PCH domains formation [22, 23]. This widely observed spatial organization of PCH domains could significantly influence transcription and other genome functions [24], such as silencing of euchromatic genes transposed near or in PCH genomic regions [25–27]. Furthermore, PCH-PCH interactions have recently been proposed to drive the global genome architecture [28].

In addition to PCH and peritelomeric heterochromatin, regions of H3K9me2/3 enrichment are also present in the euchromatic genome [29–31]. Previous studies of a large block (∼1 Mb) of *Drosophila* heterochromatin inserted in subtelomeric euchromatin (*Bw^D^*) [32, 33], revealed that large, repetitive, H3K9me2/3 and HP1a-enriched regions in the euchromatic genome can spatially interact with the main PCH domain despite their separation by a large linear distance along the chromosome. However, it remains unknown whether the more prevalent, smaller (tens of Kbs), and naturally occurring H3K9me2/3 enriched regions in the euchromatic genome (or “H3K9me2 islands”), such as those associated with epigenetically silenced transposable elements (TEs) [34, 35], also spatially contact the larger PCH domain.

We currently lack a global and in-depth understanding of the 3D organization of PCH domains, their interactions with the euchromatic genome, and the associated functional importance. To address these questions, we developed a novel method that tackles the sequence complexity of PCH to analyze Hi-C data, and used it to study the 3D organization of PCH domains. Combined with cytological analysis, we provide a comprehensive picture of the 3D structure of PCH domains in late-stage *D. melanogaster* embryos. Our analysis reveals highly heterogeneous contact frequencies among PCH regions, suggesting hierarchical ordering within the domain. Surprisingly, despite being far from PCH on linear chromosomes, euchromatic loci enriched with H3K9me2/3 can dynamically interact with the main PCH domain, and such interactions show properties consistent with liquid-liquid phase separation and influence individual fitness. Our study demonstrates that the spatial interactions among H3K9me2/3 enriched regions both in PCH and the euchromatic genome can have a fundamental impact on genome organization and, potentially, genome function.

## Results

### Hierarchical organizations of PCH domains

To decipher the 3D organization of PCH domains, we overcame technical limitations inherent to analyzing repeated DNA sequences and developed a new method that includes repetitive DNAs highly represented in PCH regions to analyze Hi-C data (Figure 1A **and Figure S1).** The Release 6 *D. melanogaster* genome is the most complete genome among all multicellular eukaryotes, and includes a nearly full assembly of the non-satellite PCH DNA [36, 37]. The genomic boundaries between PCH and euchromatin have also been epigenetically identified [31]. The annotated assembly allowed us to include three types Hi-C reads that originate from PCH DNA (Figure 1A): 1) unique single-copy sequences within PCH (e.g. protein coding genes, “unique”), 2) simple repeats known to be enriched in PCH (“repeat”, **Table S1**), and 3) sequences that map to multiple sites in the PCH (i.e. non single-locus mapping, “multi”). We used these sequence classifications to assess contact frequencies between PCH regions, and between PCH and H3K9me2/3-enriched regions in the euchromatic genome (Figure 1B and below), using published Hi-C data from 16-18hr *D. melanogaster* embryos [38].

**Figure 1.**
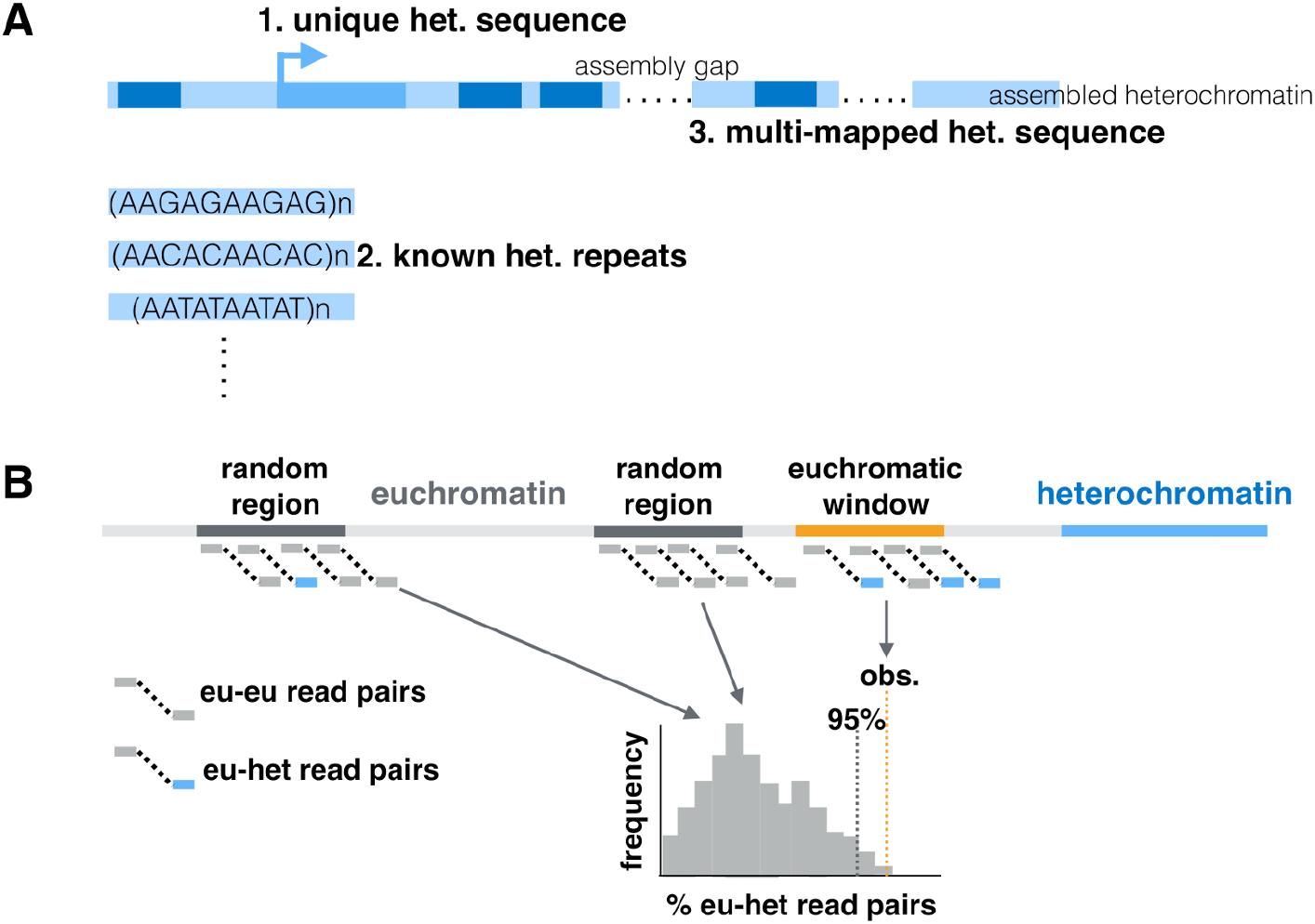
New approaches for analyzing 3D organization of PCH domains. (A) Three types of PCH-derived sequences were included in the Hi-C analysis: 1) reads mapped to single-copy sequence in the epigenetically defined PCH regions (“unique” reads, 2.4% of filtered Hi-C reads (see **Figure S1**)), 2) reads mapped to known heterochromatic simple repeats (“repeat” reads, 6.44%), or 3) reads mapped to non-unique sequences (dark blue) that are present within epigenetically defined PCH regions (“multi” reads, 3.0%). (B) Methods for assessing if a H3K9me2-enriched euchromatic region displays exceptional 3D contacts with PCH. The observed percentage of euchromatin-PCH read pairs for an H3K9me2 enriched euchromatic region is compared to a null distribution generated using randomly selected, non-H3K9me2 enriched euchromatic regions to estimate *p-value*.

Analyses of the formation and function of 3D PCH domains generally assume they are homogeneous, despite the fact that they contain coalesced PCH regions from different chromosomes that have high sequence heterogeneity. To investigate potential substructures within the PCH domains, we focused on Hi-C read pairs in which both ends mapped uniquely to PCH genomic regions (“unique” PCH reads, Figure 1A) because of their known chromosomal locations. In addition to PCH regions on the 2^nd^, 3^rd^, and X chromosomes, the entire 4^th^ and Y chromosomes were included in the analysis because the entirety of these two chromosomes are enriched with heterochromatic marks [31, 39]. We estimated the number of Hi-C read pairs coming from any two of the 100kb PCH regions. Using a sequential exclusion approach (see Methods), we identified three types of prevalent spatial interactions among PCH regions: within an arm (intra-arm), between arms of the same chromosome (inter-arm), and between arms of different chromosomes (inter-chromosome). The most frequent interactions were among PCH windows on the same chromosomal arm, which accounts for 98.08% (replicate 1, Figure 2A) and 97.15% (replicate 2, **Figure S2; and see Figure S3**) of parsed Hi-C read pairs (see **Table S2** for the number of read pairs supporting each interaction). Interactions among windows within PCH arms are stronger than PCH-euchromatin interactions on the same arm (**Figure S4 and S5**), suggesting that PCH arms (e.g. 2L PCH) are organized into distinct “territories.” Exclusion of intra-arm interactions revealed strong spatial interactions between PCH regions flanking the centromeres (inter-arm, i.e. 2L-2R, 3L-3R), which accounted for 34.72% and 35.88% (replicate 1 and 2) of the remaining read pairs (0.67% and 1.02% of total unique PCH-PCH read pairs respectively), and specific inter-chromosome interactions, mainly 3L −4 (9.68% and 9.49% of non-intra-arm read pairs). To quantitatively investigate whether these interactions are exceptional, we compared the observed percentage of read pairs against expectations that are based on either theoretical mappability [40] or empirically observed number of reads mapped to PCH on each chromosome arm (see Methods, Figure 2B) We also performed permutation tests for the latter to evaluate the statistical significance. Contact frequencies between 2L-2R, 3L-3R, and 3L-4 are indeed significantly more than expected (compared to both expectations, permutation *p-value* < 0.0001). Finally, we excluded all intra-chromosome interactions to specifically study contact frequencies between PCH regions on different chromosomes (Figure 2B). The relative frequencies of most inter-chromosome associations did not exceed expectations (e.g. 2L-3L), suggesting random contacts across cell populations. However, frequencies of 3D contacts between 3^rd^ chromosome PCH and the 4^th^ chromosome (3L-4, 3R-4) were exceptionally high (compared to both expectations, permutation *p-value* < 0.0001). Contact frequencies between 2L-4, 2R-4, and 3R-Y were also significantly more than expected.

**Figure 2.**
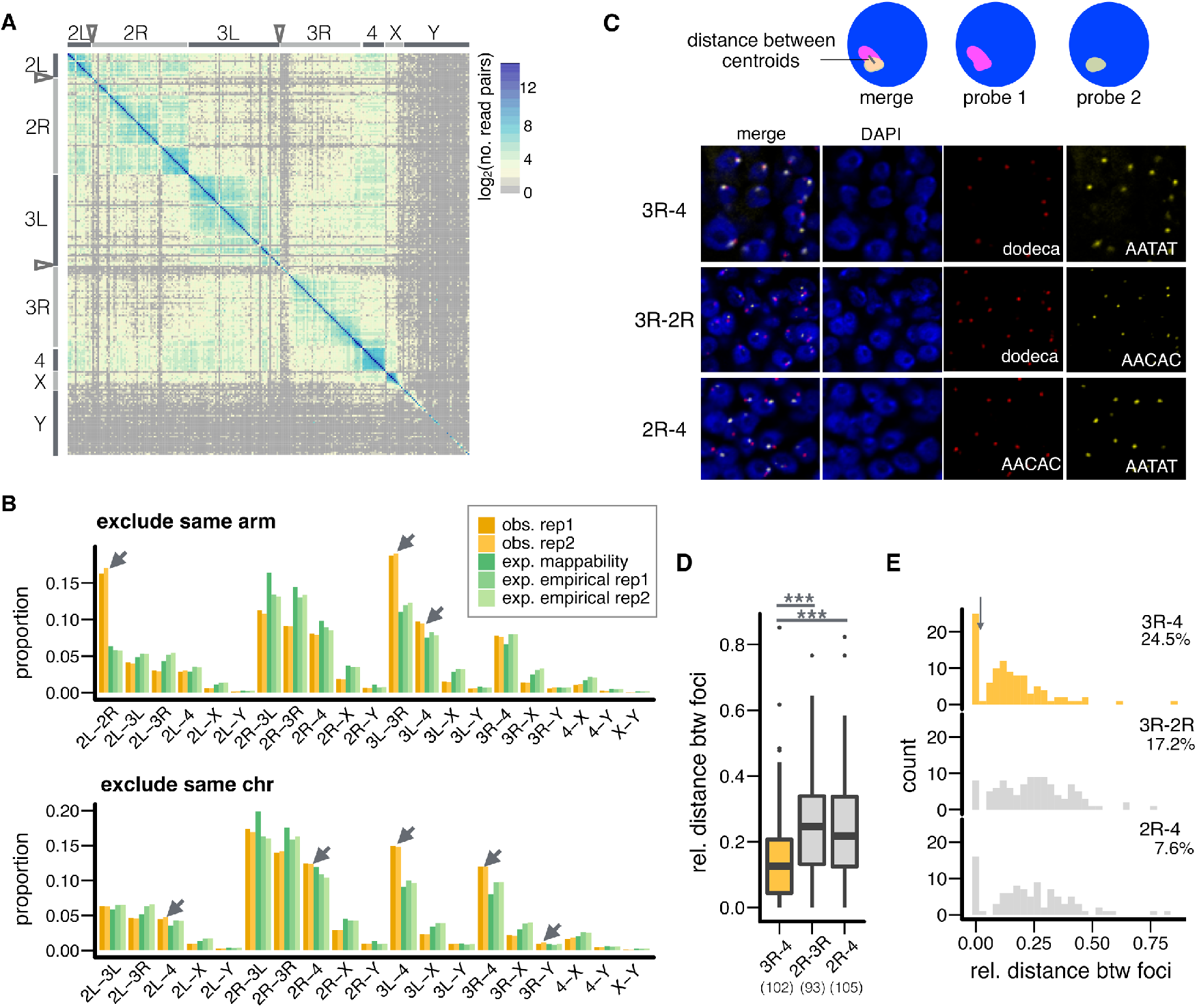
Differential spatial interactions between PCH regions on different chromosomes. **(A)** Heatmap for the number of Hi-C read pairs supporting the spatial interactions between pairs of 100kb PCH windows (total 189 windows). Replicate 1 is shown (see **Figure S2** for replicate 2). Centromeres are denoted by arrowheads and only PCH regions are shown. (B) Barplots for the observed and expected proportion of read pairs supporting spatial interactions between PCH on different chromosomes, excluding intra-arm (above) and inter-arm (below) interactions. Interactions that are more than expected and have significant permutation *p-values* (all *p* < 0.0001) are denoted with arrows. (C) Example showing how distance between foci was estimated (top) and representative images of embryonic cells stained with DAPI (DNA, blue) and FISH probes recognizing indicated PCH regions (3R-4, 2R-3R, and 2R-4, pink and yellow) (bottom). (D,E) Boxplot (D) and histogram (E) showing the relative distance between PCH foci. Orange box/bars are for exceptional PCH interactions (3R-4) while gray ones are for other interactions. In (D), numbers of nuclei counted are in parentheses. Center lines: median, box limits: upper and lower quartile. Points: outliers. In (E), threshold for nuclei with overlapping foci is denoted with arrow, and the percentages denote nuclei with overlapping foci. FISH analysis based on absolute distance led to the same conclusions (**Figure S6**). ** *p <* 0.01, *** *p <* 0.001.

The spatial interactions detected with Hi-C represent a superimposition of different chromosome conformations within cell populations. To investigate the prevalence and cell- to-cell variability of identified 3D interactions, we performed single-cell fluorescence in situ hybridization (FISH) on embryos of the same genotype and stage as those used for Hi-C. In *D. melanogaster,* different simple repeats are specifically enriched in the PCH regions of certain chromosomes [41]. This allowed us to ask if chromosome-specific probes that label simple repeats from PCH regions that displayed exceptional Hi-C spatial interactions (e.g. 3R-4) colocalized more often than probes from the same chromosomes with lower frequency interactions (2R-3R and 2R-4). We measured the “relative distance,” defined as the distance between FISH signal centroids divided by the nuclear radius (Figure 2C), to account for variable cell size at late embryonic stages. The relative distance between 3R (dodeca)-4^th^ chromosome (AATAT) is significantly shorter than 2R (AACAC)-3R or 2R-4 (*Mann-Whitney test, p* = 0.0001 (3R-4 vs 2R-3R) and <10^-6^ (3R-4 vs 2R-4), Figure 2D). For all three pairs of interactions, the distribution of relative distance is bimodal (Figure 2E), with a sharp peak near zero. We defined two foci as ‘overlapping’ when their distances were shorter than this natural threshold (denoted by arrow in Figure 2E). Consistent with the Hi-C results, the proportion of nuclei with overlapping foci was higher for 3R-4 than for 2R-3R or 2R-4 (*Fisher’s Exact test, p* = 0.22 and 0.0006 respectively, Figure 2E). Overall, both Hi-C and FISH analyses demonstrate a hierarchical 3D organization of PCH domains.

### Euchromatic regions enriched for H3K9me2 show 3D contacts with PCH

The coalescence of PCH regions and large blocks of translocated heterochromatin in the euchromatic genome (e.g. *Bw^D^,* [32, 33]), as well as the observations of the formation of HP1a liquid droplets both *in vitro* and *in vivo* [22, 23], led us to predict that small regions enriched for H3K9me2/3 and HP1a in the euchromatic genome could also spatially associate with the main PCH domains. To test this hypothesis, we identified euchromatin-PCH Hi-C read pairs, which contain sequences from single-copy, euchromatic regions paired with *any* PCH sequence (i.e. all three categories of PCH sequences, Figure 1A). We then estimated, among Hi-C read pairs whose one end mapped uniquely to a specific euchromatic region, the percentage of euchromatin-PCH read pairs (Figure 1B). We generated null distributions for the percentage of euchromatin-PCH Hi-C read pairs using random non-H3K9me2/3 enriched euchromatic regions to calculate empirical *p-values* (Figure 1B). Euchromatic regions with exceptional percentage of euchromatin-PCH Hi-C read pairs (empirical *p-values* < 0.05) were considered to interact spatially with PCH (see Methods).

We identified by ChIP-seq 496 H3K9me2-enriched regions (defined as “H3K9me2 islands,” 290bp - 21.63Kb, with an average size of 3.84 kb) in the euchromatic genome (>0.5 Mb distal from the epigenetically defined euchromatin-PCH boundaries) in embryos of the same genotype and stage as the Hi-C data (see Methods). Of these H3K9me2 islands, 13.91% (n = 69) and 8.67% (n = 43) displayed significant spatial associations with PCH in either or both Hi-C replicates, respectively (Figure 3A). These numbers are significantly higher than expected (i.e. 5% of the H3K9me2 islands would be significant under null expectation; *binomial test, p* = 0.00059 (both) and 3.04×10^-14^ (either)). Thus, we conclude that H3K9me2 islands are more likely to spatially interact with PCH than euchromatic regions without H3K9me2 enrichment in the euchromatic genome. For subsequent analyses, we focused on H3K9me2 islands that significantly interacted with PCH in *both* Hi-C replicates (hereafter referred to as “EU-PCH” associations).

**Figure 3.**
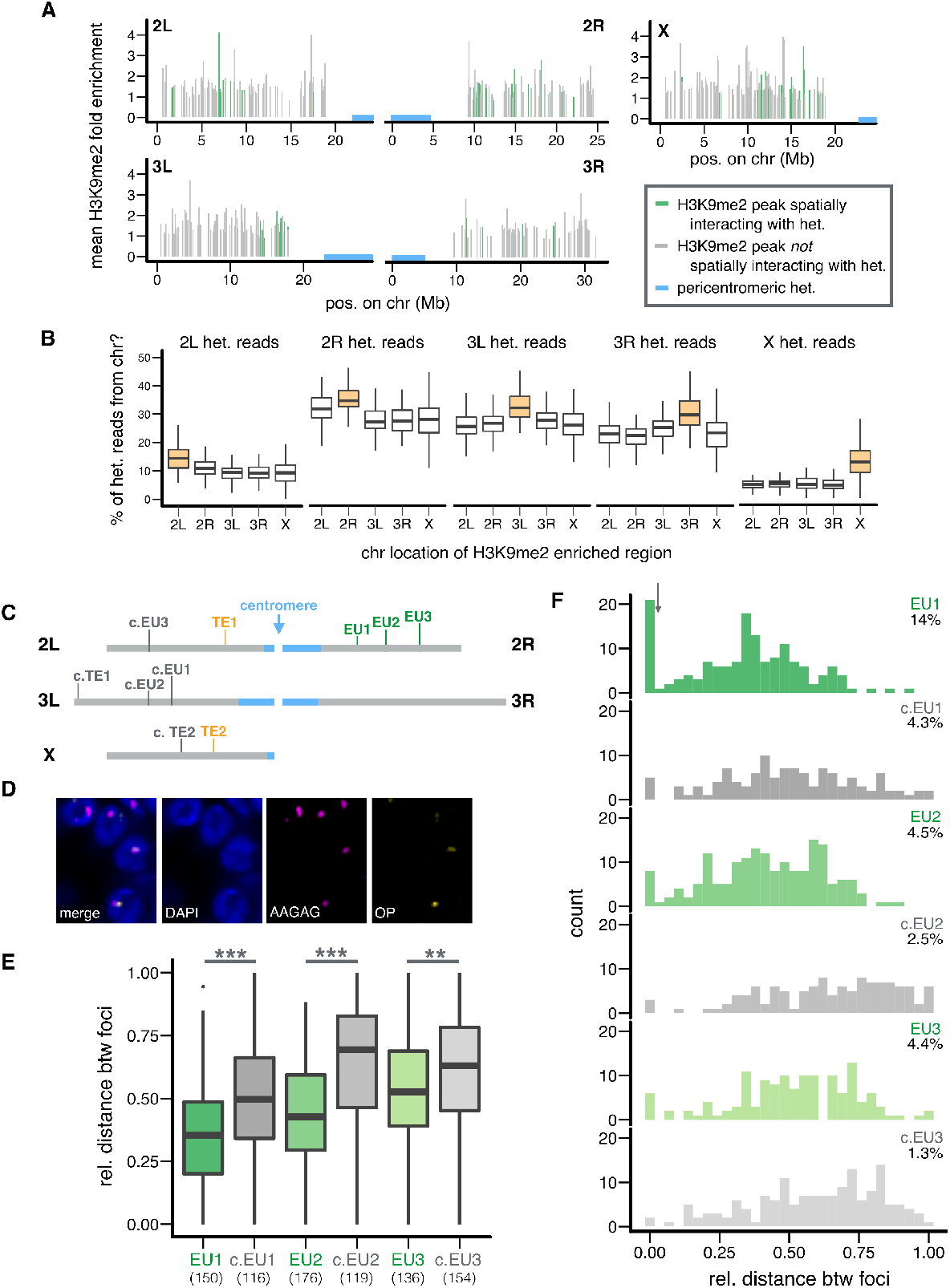
H3K9me2 islands are in 3D contacts with PCH domains. (A) Genomic distribution and average H3K9me2 enrichment level of H3K9me2 islands with (green) or without (gray) 3D interactions with PCH (blue). (B) The percentage of Hi-C reads coming from PCH regions on a particular chromosome (y-axis) is compared between H3K9me2 islands on the same (orange) or other (white) chromosomes. Replicate 1 is shown and see **Figure S8** for replicate 2. (C) Locations of H3K9me2 islands and TEs chosen for FISH analysis. Euchromatin (gray), PCH (blue). (D) Representative image of nuclei stained with DAPI (DNA, blue) and FISH probes for EU1 (OP=Oligopaint probe, yellow) and PCH (AAGAG, pink). Also see **Figure S10**. (E,F) Boxplot (E) and histogram (F) showing the relative 3D distance between PCH and indicated euchromatic regions (with PCH interaction = green, without = gray). In (E), numbers of nuclei counted are in parentheses. In (F), threshold for nuclei with overlapping foci is denoted with arrow, and the percentages denote nuclei with overlapping foci. FISH analysis based on absolute distance led to the same conclusions (**Figure S11**). In (B, E) Center lines: median, box limits: upper and lower quartile. Points: outliers. ** *p <* 0.01, *** *p <* 0.001.

We found that H3K9me2 islands with PCH interactions have shorter linear distance to PCH regions along the chromosome compared to H3K9me2 islands that lacked PCH interactions (*Mann-Whitney U test, p* < 10^-4^, **Figure S7**), suggesting that proximity to PCH on a linear chromosome is a strong defining feature for the tendency to spatially interact with PCH. For each H3K9me2 island, we calculated the percentage of unique PCH reads from each chromosome arm (e.g. percentage of EU-2L PCH read pairs). For PCH region on a particular arm, H3K9me2 islands on the very same arm always have the highest such percentage (e.g. 2L euchromatic regions have the highest percentage of EU-2L PCH read pairs), followed by those on the other arm of the same chromosome (Figure 3B and **Figure S8**). This echoes the observed strong tendency of “intra-arm” PCH-PCH interactions, followed by “inter-arm” PCH-PCH interactions (Figure 2A **and 2B**).

Interestingly, H3K9me2 islands that show spatial interactions with PCH have higher fractions of coding sequences when compared to H3K9me2 islands without PCH interactions (*Mann-Whitney U test, p* = 0.0015, median: 70.1% (with) and 30.4% (without)). In addition, these regions are more likely located within active Topologically Associated Domains (TADs) identified at the same embryonic stage [8] than H3K9me2 islands without PCH interactions (*Fisher’s Exact Test, p* = 0.0078, **Table S3)**. Using previously reported segmentations of the *D. melanogaster* genome into combinatorial chromatin states [42, 43], we also found that significant EU-PCH contacts are more likely to involve euchromatic regions in active states: Red or Yellow chromatin (*Fisher’s Exact test, p* = 0.021), or modEncode State 1-4 (*p* < 10^-4^ (S2) and =0.011 (BG3), **Table S3**). These regions are also depleted for chromatin states that lack obvious enrichment for histone modifications and/or protein binding: “null” TADS (*Fisher’s Exact test, p* = 0.03), black chromatin (*p* < 10^-3^), and modEncode State 9 (*p =* 0.008 (S2), **Table S3**). It is currently unclear why PCH associations would be enhanced for H3K9me2 islands containing coding genes or active chromatin marks. It is worth noting that PCH associations were not correlated with the following properties of H3K9me2 islands: autosome or sex chromosome linkage (*Fisher’s Exact test, p* = 0.27), size of the enriched region (*Mann-Whitney U test, p* = 0.31), or the average level of H3K9me2 enrichment (*Mann-Whitney U test, p* = 0.91). Analysis of significant EU-PCH interactions in *either* replicate reached the same conclusions (**Table S4**).

To validate the EU-PCH 3D interactions identified by Hi-C analysis, we performed FISH using Oligopaint probes [44–46] targeting 30.5-42.9kb euchromatic regions (**Table S5**) and probes that broadly mark PCH (AAGAG, a satellite enriched in PCH regions of all chromosomes, [47, 48]). We focused on three 2R windows covering H3K9me2 islands that spatially interact with PCH (EU1-3). Because we observed that the linear distance to PCH genomic regions is a strong predictor for whether a H3K9me2 island interacts with PCH (see above), for each of these regions, we chose a matching “control” window that is at a similar linear distance from PCH genomic regions and does not have H3K9me2 enrichment (c.EU1-3, see Figure 3C for genomic locations of chosen regions, see **Figure S9** for their H3K9me2 enrichment level, and Figure 3D and **Figure S10** for representative cell images). Consistently, we observed that H3K9me2 islands displaying PCH interactions in the Hi-C analysis are closer to PCH in 3D space than linearly equidistant euchromatic regions that lack H3K9me2 enrichment (*Mann-Whitney U test, p* < 10^-6^ (EU1 vs c.EU1), < 10^-13^ (EU2 vs c.EU2), and 0.0025 (EU3 vs c.EU3), Figure 3E), confirming the observations made by Hi-C analysis. This difference is also reflected in the higher proportion of cells in which the two foci overlap compared to the control regions (Figure 3F). It is worth noting that the comparatively lower frequency of overlapping foci for EU2 and EU3, when compared to EU1, could result from the fact that these two regions are much farther from the PCH, and thus less likely to spatially interact with PCH than EU1 (see above). This could potentially lead to lower statistical power and thus the comparison of proportion of overlapping foci between focused and control regions is only statistically significant for EU1 (*Fisher’s Exact test, p* = 0.007 (EU1 vs c.EU1), 0.37 (EU2 vs c.EU2), and 0.15 (EU3 vs c.EU3)). Overall, the Hi-C and FISH analyses reveal that even short stretches of H3K9me2-enrichment in the euchromatic genome can coalescence with the main PCH domains. We would like to note that the focused regions (EU1-3) and control regions (c.EU1-3), though are of similar distance to PCH, are not on the same chromosome, and biases that we were unaware of could have led to the observed results. Stronger evidence would come from comparing the 3D organization of homologous sequences with and without H3K9me2 enrichment (see below).

### 3D PCH contacts include euchromatic TEs enriched for H3K9me2

Naturally occurring TE insertions in the euchromatic genome can acquire H3K9me2/3 marks that often extend into flanking regions, including genes [34,35,49,50], and we predict that these could also spatially contact the main PCH domains. While non-TE induced H3K9me2/3 enriched regions in the euchromatic genome are commonly *shared* between individuals (e.g. **Figure S9**), most TE insertions are polymorphic (i.e. not present in all individuals) in the *Drosophila* population [51–53], leading to varying H3K9me2 enrichment between individuals and strains (e.g. **Figure S12**, [35]). Accordingly, we compared the H3K9me2 enrichment level around euchromatic TE insertions in the strain used for Hi-C (ORw1118) with that of homologous sequences in strains without the respective TEs (wildtype) to identify TE-induced H3K9me2 islands, as performed previously [35]. This approach identifies H3K9me2 enrichments that are broad and/or low in enrichment level, and therefore often missed by custom pipelines that rely on identifying “sharp peaks” (reviewed in [54, 55]). Our analyses were restricted to 106 TEs that displayed H3K9me2 spreading into at least 1kb of flanking DNA (65% of identified TEs in strain ORw1118, see Methods), with an average of 4kb and maximum of 18kb of H3K9me2 spread. Among these TEs, 13.21% (n = 14) and 7.55% (n = 8) displayed significant spatial interactions with PCH (*p* < 0.05) in either or both Hi-C replicates respectively (see **Figure S13** for their genomic distribution), which is significantly more than expected (*binomial test, p* = 8.38×10^-4^ (either) and 0.26 (both)). As a contrast, only 1.75% of TEs without H3K9me2 enrichment (n = 1) display PCH interactions. We focused on analyzing the 14 TEs showing significant PCH-contact in *either* replicate, while analyses restricted to eight TEs significant for *both* replicates was qualitatively similar (**Table S6**). Similar to non-TE induced H3K9me2 islands, TEs spatially interacting with PCH are closer to PCH genomic regions on the linear chromosome than those that do not interact with PCH (*Mann-Whitney U test, p* = 0.037, **Figure S14**). PCH-interacting TEs include those from *roo, pogo, 17.6, mdg3, FB,* and *S* families. However, they were not significantly enriched for any specific TE family (*Fisher’s Exact Test* for individual TE family*, p* > 0.26), class, type, or sex-chromosome linkage (**Table S6**).

The polymorphic nature of TEs offers a rare opportunity to compare the 3D conformations of homologous sequences with and without TE-induced H3K9me2/3 enrichment. To validate the Hi-C results, we performed FISH analysis focusing on two TEs that are present in the Hi-C strain (ORw1118) but absent in another wildtype strain. These two TEs also induced ORw1118-specific enrichment of H3K9me2 (**Figure S12**) and spatially interact with PCH (TE1-2, Figure 3C). As controls, we included two additional ORw1118-specific TEs that did not interact with PCH and do not have H3K9me2 enrichment (c.TE1-2, Figure 3C and **Figure S12**). Our FISH used Oligopaint probes that target *unique regions* flanking the selected euchromatic TE insertions (**Table S5**) and probes that broadly mark PCH (see **Figure S10** for representative cell images). For TE1 and TE2, the relative 3D distance to PCH signals is shorter in ORw1118 than in wildtype (*Mann-Whitney U test, p* = 0.0004 (TE1) and *p* = 0.015 (TE2), Figure 4A). Interestingly, the distribution of relative distance between TE1/TE2 and PCH is bimodal for ORw1118 nuclei but unimodal for wildtype, which lacks the peaks around zero, or nuclei with overlapping foci (Figure 4B). Indeed, there are more nuclei with overlapping foci in ORw1118 than in the wildtype (*Fisher’s Exact Test, p* = 0.0003 (TE1) and 0.070 (TE2)). Importantly, these between-strain differences were not observed for control TEs that lacked PCH interactions (*Mann-Whitney U test, p* = 0.55 (c.TE1) and 0.91 (c.TE2), *Fisher’s Exact test, p* = 0.49 (c.TE1) and 1 (c.TE2), Figure 4A **and 4B**). This comparison of *homologous* regions with and without euchromatic TEs suggests that H3K9me2 enrichment is required for spatial contacts between euchromatic regions and PCH domains.

**Figure 4.**
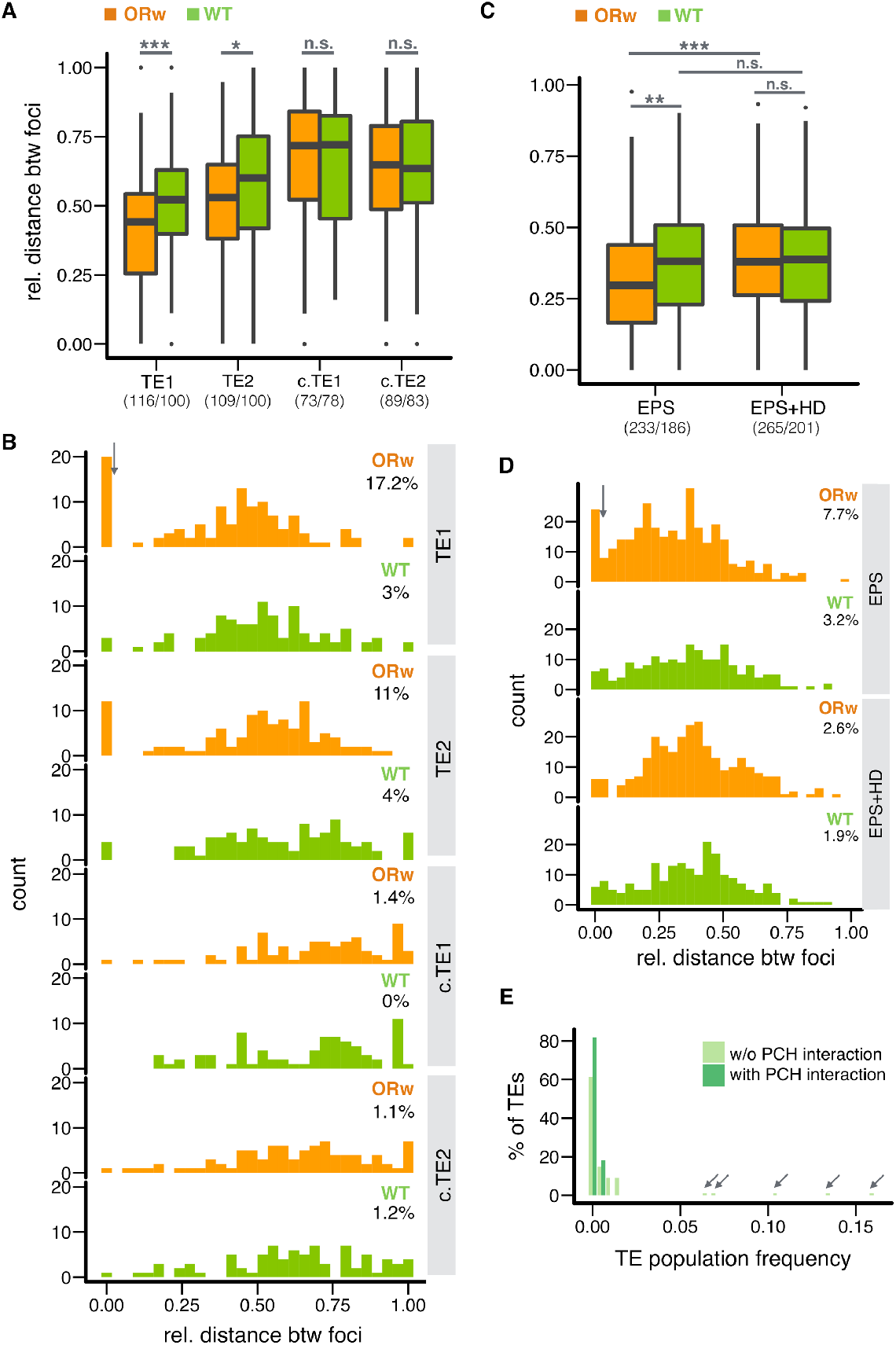
Euchromatic TEs show 3D contacts with PCH, and such interactions are sensitive to 1,6-hexanediol. (A, B) Boxplot (A) and histogram (B) showing the relative distance between euchromatic TE neighborhood and PCH. Relative distance was estimated for ORw1118 (ORw, orange, TE present) and wild type (WT, green, no TE present) embryonic cells. (C, D) Boxplot (C) and (D) histogram comparing TE1-PCH relative distance between genotypes and between treatments. Relative distance was estimated for permeabilized ORw and WT embryos (EPS, see Methods) and permeabilized ORw and WT embryos with 1,6-hexanediol treatments (EPS+HD). In (A, C), numbers of nuclei counted are in parentheses. In (B, D), threshold for nuclei with overlapping foci is denoted with arrow, and the percentages denote nuclei with overlapping foci. FISH analysis based on absolute distance led to the same conclusions (**Figure S15 and S17**). (E) Population frequencies of TEs with and without PCH interaction. Note that high frequency TE insertions (population frequency > 0.05, arrows) all show no PCH interactions. In (A, C), Center lines: median, box limits: upper and lower quartile. Points: outliers. * *p* < 0.05, ** *p <* 0.01, *** *p <* 0.001, n.s *p >* 0.05.

### Euchromatin-PCH 3D contact is sensitive to perturbing liquid-liquid phase separation

The coalescence of PCH regions located on different chromosomes into 3D PCH domains in *Drosophila* exhibits properties characteristic of liquid-liquid phase separation, including sensitivity to 1,6-hexanediol [23], a mild perturbant of hydrophobic interactions [56]. To investigate if the 3D contacts between H3K9me2 islands and PCH domains is mediated by similar biophysical interactions, we used FISH to compare the 3D distance between PCH and H3K9me2 islands that displayed significant PCH interactions (see above) in permeabilized embryos with and without 1,6-hexanediol treatment (see Methods). We focused on TE1 because it is ORw1118-specific and leads to strain-specific H3K9me2 enrichment. This allows comparisons between genotypes with and without TEs to investigate whether the sensitivity to 1,6-hexandiol treatment is H3K9me2-enrichment dependent (see Methods, **Figure S16**). We observed significantly longer TE1-PCH relative 3D distance (orange in Figure 4C, *Mann-Whitney test, p* < 10^-4^) and fewer nuclei with overlapping foci (orange in Figure 4D, *Fisher’s Exact test, p* = 0.02) in ORw1118 embryos treated with 1,6-hexanediol compared to untreated controls. In contrast, no such difference was observed in wildtype embryos, which do not have the TE insertion and thus no frequent TE1-PCH 3D contacts (green in Figure 4C **and 4D**, *Mann-Whitney test, p* = 0.74, and *Fisher’s Exact test, p* = 1). Importantly, the significant difference in TE1-PCH 3D distance between genotypes with and without TE insertion is only observed for embryos *without* 1,6-hexanediol treatments (*Mann-Whitney test, p* = 0.0037, *Fisher’s Exact test, p* = 0.057), but not for those *with* the treatment (*Mann-Whitney test, p* = 0.77 and *Fisher’s Exact test, p* = 0.55, Figure 4C **and 4D**). The sensitivity of TE-PCH 3D contacts to 1,6-hexanediol is consistent with the spatial interactions between H3K9me2 islands and PCH domains being mediated by liquid fusions, an emergent property of liquid-liquid phase separation.

### Euchromatin-PCH 3D contact may influence individual fitness

A dominant factor governing the population frequencies of TEs (presence/absence in a population) is natural selection against their deleterious fitness impacts [51,57,58]. We estimated the population frequencies of studied TE insertions (in ORw1118 genome) in a large panmictic African population ([59], see Methods). TEs with PCH interactions have significantly lower mean population frequencies than TEs without (*t-test, p* = 0.0042, mean frequency 9.7×10^-4^ (with spatial interaction) and 9.6×10^-3^(without), see Methods) and their frequency spectrum is more skewed towards rare variants (Figure 4E). Both of these observations support stronger selection against TEs with PCH interactions than other TEs [51,57,58], which could result from the negative functional consequences of TE-PCH 3D interactions. It is worth noting that even 0.01% variation in fitness, which could be rarely detected in a laboratory, can result in large differences in population frequencies in nature.

Multiple other factors have been correlated with TE population frequencies, such as TE type, chromosome linkage, and recombination rate [52, 60], and could also contribute to the low population frequencies of TEs displaying PCH interactions. However, TEs with and without PCH interactions do not differ in their class, type, chromosome linkage (**Table S6**) or local recombination rate (*Mann-Whitney U test, p* = 0.40). On the other hand, we did observe that TEs with PCH interactions tend to be closer to genes than TEs without such interactions (*Mann-Whitney U test, p* = 0.065). The stronger selection against TEs with PCH interactions could thus result from either the direct functional impact of PCH spatial contacts on adjacent genes (see Discussion) and/or other TE-mediated functional impacts along the linear chromosome (such as disrupting regulatory non-coding sequences).

## Discussion

An appreciable fraction of most eukaryotic genomes comprises constitutive heterochromatin, which is enriched for megabases of repetitive DNA and predominantly localizes around centromeres (PCH). However, because of technical difficulties associated with repetitive DNA, we have lacked a global and in-depth understanding of the 3D organization of the PCH domain, which encompasses at least a fifth of the human [61] and a third of the *D. melanogaster* genomes [37]. In this study, we aimed to provide a comprehensive and detailed picture of the 3D organization of PCH domains in *D. melanogaster* by combining genome-wide Hi-C analyses and cytological FISH studies. We developed a novel analysis approach that overcomes the challenges posed by repeated DNAs when determining 3D contact frequencies from Hi-C reads. Specifically, we relaxed the single-locus mapping restriction to include reads originating from the abundant repetitive DNA in PCH, and used different combinations of PCH reads (single-locus mapping or not) depending on the question being addressed. Our investigations reveal significant, new insights into the interactions between different PCH regions and their 3D contacts with the euchromatic genome.

The coalescence of PCHs on different *D. melanogaster* chromosomes contributes to the formation of a large PCH domain in 3D nuclear space. However, we found that DNA contacts within the PCH domain are far from homogeneous. Our Hi-C analysis revealed the strongest interactions (∼98%) involve PHC regions on the same chromosome arm (e.g. 2L), suggesting PCH regions from each arm are organized into distinct “territories” (Figure 5). This is similar to identified chromosome territories for the euchromatic genome [6,8,62– 64]. PCH regions from all the chromosomes do interact. However, some interactions occur more often than random, in particular the inter-arm (2L-2R, 3L-3R) and specific inter-chromosomal (3L/3R-4) 3D associations. Most strikingly, ∼14% of identified H3K9me2-enriched regions in epigenomically-defined euchromatin display preferential 3D contacts with the central PCH domains (Figure 3A and 4A). Importantly, quantitative FISH analysis provides cytogenetic support for the Hi-C results. The bimodal distributions of PCH-PCH or EU-PCH distances in nuclei (Figure 2F, 3G, 3E) also demonstrate that these 3D contacts are dynamic and can vary among cells, as previously shown for euchromatic *Hox* loci in mouse [65]. Importantly, polymorphic TE insertions in euchromatin allowed us to directly compare homologous sequences with and without H3K9me2 enrichment, which strongly supports the conclusion that H3K9me2 enrichment is required for EU-PCH 3D contacts.

**Figure 5.**
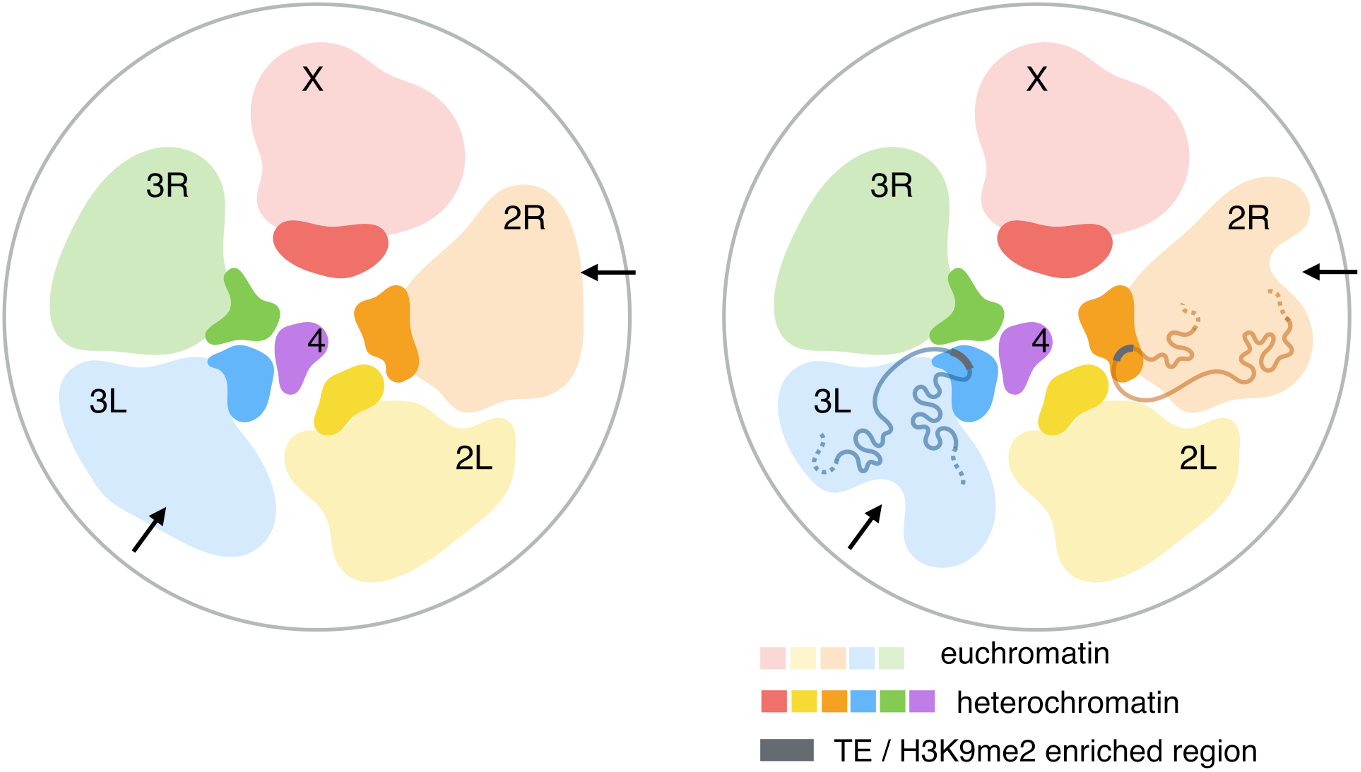
Proposed spatial architecture of *D. melanogaster* genome. PCH genomic regions located on different chromosomes coalesce to form the 3D PCH domains, or chromocenters. PCH regions (darker color) and the euchromatic genome (lighter color) form their own separate territories. PCH regions on different chromosomes interact, with inter-arm (2L-2R, 3L-3R) and inter-chromosomal 3^rd^-4^th^ chromosome 3D interactions being more frequent than random expectations. 3D contacts between polymorphic H3K9me2/3-enriched regions in the euchromatic genome (gray bar) lead to varying 3D genome conformations between individuals (arrows). 3D structures of the euchromatic genome were based on [66].

Overall, the Hi-C and FISH analyses reveal a previously unknown picture of the 3D architecture of the PCH domains (Figure 5): the spatial interactions within the domains, instead of being random, are hierarchical. In addition, despite the separation of euchromatic and PCH territories on the same chromosome arm [66], short stretches of H3K9me2/3 enrichment in the euchromatic genome (with and without TEs) also dynamically interact with the main PCH domains. Both PCH-PCH and EU-PCH interactions happen most often within chromosome arms, which is consistent with the predictions of polymer physics on chromosome folding [67, 68]. Specific spatial contacts between PCH regions located on different chromosomes are surprising, but nevertheless consistent with the observed coalescence of PCH of all chromosomes into chromocenters. The varying frequencies of inter-chromosomal interactions could result from non-random positioning of PCH regions upon mitotic exit [32]. Alternatively, variation in biophysical properties (e.g. viscosity or varying protein compositions [69]) among PCH domains arising from specific chromosomes could result in different frequencies of liquid-liquid fusion. Finally, the tendency of H3K9me2 islands to interact with PCH strongly depends on the distance to PCH on a linear chromosome. This suggests that euchromatic regions and PCH could be in transient proximities with a frequency that largely follows polymer physics of chromosome folding. The enrichment of H3K9me2/3 and the reader protein HP1a at specific euchromatic loci would then allow their liquid-like fusion with HP1a-enriched PCH, resulting in frequent and/or maintained EU-PCH 3D interactions.

Importantly, the population genetic analysis reveals that euchromatic TEs with PCH interactions have lower population frequencies than TEs lacking frequent PCH contacts (Figure 4E), suggesting that EU-PCH 3D interactions may influence individual fitness. What are the potential functional consequences of TE-PCH interactions that could influence individual fitness? TE-PCH interactions could lead to increased TE-induced enrichment of repressive epigenetic marks on neighboring sequences/genes. However, we found no difference in the extent or the magnitude of H3K9me2 spread around TEs with and without PCH interactions (*Mann-Whitney U test, p* = 0.30 (extent) and 0.53 (magnitude), **Figure S18**), suggesting that TE-PCH interactions influence other aspects of nuclear organization critical for gene regulation and/or other genome functions. For instance, 3D interactions between PCH and TEs could bring neighboring euchromatic genes into the PCH domains and result in aberrant silencing. On the other hand, the enrichment of HP1a, and likely spatial localization in the PCH domains, can play positive roles for the expression of genes in both PCH [24,70,71] and the euchromatic genome [72–74]. Still another possibility is that the spatial contact with PCH on one chromosome may “drag” its homolog to the same nuclear compartment due to somatic homolog pairing (reviewed in [75]), resulting in *trans*-silencing [76]. A preliminary analysis found that ∼15% of heterozygous TEs induced H3K9me2 enrichment not only *in cis,* but also *in trans* on the homologous chromosome without the TE insertion (i.e. *trans-*epigenetic effects, **Supplementary Text**). Accordingly, the fitness consequences of TE-PCH spatial interactions could potentially result from their positive as well as negative impacts on the expression of genes *in cis* or *in trans* to TEs, or from influencing other genome functions, such as replication and repair.

It is important to note that TEs comprise an appreciable fraction of euchromatic genomes in virtually all eukaryotes [77]. For instance, more than 50% of assembled human euchromatin contains TEs or TE-derived sequences [61, 78], many of which are interspersed with actively transcribed genes and can influence gene expression through H3K9me2/3 spreading [49]. Moreover, the presence of many TE insertions at specific locations are polymorphic between individuals in natural populations (e.g. human [79, 80], *Caenorhabditis* [81, 82], *Drosophila* [52,53,83], and *Arabidopsis* [84–86]). Spatial interactions between euchromatic TEs and PCH can thus generate polymorphic 3D organization of euchromatic genomes (Figure 5), leading to variation in critical biological functions that depend on chromosome conformations and even varying fitness between individuals. Our investigation of the spatial architecture of PCH domains could thus have strong implications for how such 3D organizations could influence gene regulation, genome function, and even genome evolution of both heterochromatin and the gene-rich euchromatin.

## Methods

### Fly strains and culture

Three *D. melanogaster* strains were used: Oregon-R w1118 (abbreviated ORw1118, [8]) and two wildtype strains, RAL315 (Bloomington Drosophila Stock Center (BDSC) 25181) and RAL360 (BDSC 25186). The latter two are part of a large collection of genomically sequenced natural *D. melanogaster* strains [87], whose TE insertion positions were previously identified [83]. Flies were reared on standard medium at 25°C with 12hr light/12hr dark cycle.

### Euchromatin-heterochromatin boundaries

To identify Hi-C reads coming from PCH genomic regions, we used epigenetically defined euchromatin-heterochromatin boundary in [31] and converted those to Release 6 coordinates using liftover (https://genome.ucsc.edu). For defining H3K9me2-enriched euchromatic regions and euchromatic TE insertions, we used 0.5 Mb inward (distal to PCH) of the epigenetically defined euchromatin-heterochromatin boundary to be conservative. The entirety of 4^th^ and Y chromosomes are enriched with heterochromatic marks [31, 39] and are considered to be entirely heterochromatic.

### Generation and analysis of H3K9me2 ChIP-seq data

We performed ChIP-seq using antibody targeting H3K9me2 (Abcam 1220) on 16-18hr embryos of ORw1118 and two wildtype strains (see above). Embryo collections and ChIP-seq experiments were performed following [35], except that sequencing libraries were prepared using NEBNext Ultra DNA Library Prep Kit for Illumina (NEB cat#E7370L) following manufacturer’s protocol and sequenced on Illumina Hi-Seq 4000 with 100bp paired-end reads. Each sample has two ChIP replicates (biological replicates) with matching inputs.

Raw reads were processed with trim_galore [88] to remove adaptors, low quality bases, and single-end reads. Processed reads were mapped to release 6 *D. melanogaster* genome with bwa mem with default parameters. Reads with mapping quality lower than 30 were removed using samtools [89]. To have enough noise for the IDR analysis (see below), we ran Macs2 [90] using broad-peak and pair-end mode, and a liberal *p-value* threshold (0.5). This was followed by performing Irreproducible Rate (IDR) analysis [91] to identify H3K9me2 enriched regions that are consistent between replicates. We defined H3K9me2-enriched regions as those with low IDR (IDR < 0.01). IDR plots for replicates for three ChIP-seq samples can be found in **Figure S19-21**.

### Identification and analysis of TE insertions

#### TEs in wildtype strains

All potential TE insertions in RAL315 and RAL360 strains were previously identified using TIDAL [83]. We used the recommended coverage ratio (read number supporting TE presence/TE absence, coverage ratio at least three) to identify TEs with high confidence in these two wildtype strains. TEs in wildtype strains are used to identify ORw1118-specific TEs (see below).

#### Identification of TEs in ORw1118

To identify TEs in the ORw1118 strain, we performed genomic sequencing. Genomic DNA was prepared from 100 ORw1118 adult female flies for each biological replicate (three biological replicates in total) with Gentra Puregene Cell kit (Qiagen cat#158388) according to the manufacturer’s instructions. Whole genome sequencing was done with overlapping 165bp pair-end Illumina sequencing on 230-240bp size genomic fragments.

We combined all three replicates of ORw1118 genomic sequencing to call TEs and quality filtered reads with Trim_galore. We identified TEs in ORw1118 also using TIDAL [83], which calls TEs with split-read methods and requires input reads to have the same length. Accordingly, we used two approaches to generate single-end reads from the original pair-end data (1) treating pair-end reads as single-end and (2) use SeqPrep (https://github.com/jstjohn/SeqPrep) to merge overlapping reads and trimmed reads to 200bp. We used the same TIDAL parameters (default) and coverage ratio (at least three) as those used in calling TEs in wildtype strains [83]. 249 called TEs overlap between the two approaches (89.2% and 89.9% of the called TEs respectively). We further removed TEs in shared H3K9me2-enriched euchromatic regions of wildtype strains (see above) or shared with wildtype strains, with the idea that local enrichment of H3K9me2 in ORw1118 cannot be unambiguously attributed to the presence of TE insertions. In total, 166 euchromatic TEs in ORw1118 were identified with these criteria.

To identify TE-induced local enrichment of H3K9me2, we used methods described in [35], which leverages between strain differences to identify TE-induced H3K9me2 enrichment regions with any shape, which oftentimes do not resemble peaks (e.g. **Figure S12**). This approach is more sensitive than other custom pipelines, which look for enrichment with “peak” shape, followed by *ad hoc* merging of sharp peaks to generate “broad peak” calls (reviewed in [54, 55]). We compared the enrichment of H3K9me2 in euchromatic TE neighborhoods in ORw1118 against wildtypes strains to estimate (1) the extent of TE-induced H3K9me2 enrichment (in kb) and (2) % of increase of H3K9me2 enrichment. We identified 106 ORw1118 TEs leading to at least 1kb spread of H3K9me2, with only 13 of them overlap with H3K9me2 enriched regions identified by Macs2.

We used the same approach as in [35] to estimate the population frequencies of ORw1118 TEs in an African population [59]. Similar to previously reported low population frequencies of TEs in *Drosophila* [51–53], only 36.36% of the 106 euchromatic TEs that induced H3K9me2 enrichment are present in a large African population [59] (i.e. 63.64% of those TEs are unique to ORw1118). This generally low population frequency of TEs is expected to limit the statistical power of comparison between TEs with and without PCH interactions. Indeed, we found that the median population frequencies for both TEs with and without PCH interactions are zero and not significantly different (*Mann-Whitney U test, p =* 0.10). Accordingly, we instead investigated whether the mean of their population frequencies differ (see main text).

### Analysis of Hi-C data

Raw Hi-C reads (two biological replicates) from [38] were downloaded from GEO and quality filtered with trim_galore. TEs are abundant in both euchromatin and heterochromatin in *Drosophila* [37, 92], and we were unable to unambiguously define which genomic compartment a TE-mapping read is from. Accordingly, we filtered reads that mapped to canonical TEs using bwa [93] and samtools [89]. Filtered reads were then mapped to release 6 *D. melanogaster* reference genome using bwa with default parameters. Three types of reads are defined as from heterochromatin. (1) Reads that uniquely mapped (mapping quality at least 30) within epigenetically defined PCH regions. (2) Reads mapped to known heterochromatic repeats (**Table S1**). (3) Reads that mapped to epigenetically defined PCH but have mapping quality equals zero, which bwa assigns to multiple-mapped reads. Mapping locations of unique PCH reads are recorded and used for both PCH-PCH and PCH-EU analysis. Other two types of PCH reads were only used for PCH-EU analysis and their mapping locations, which are multiple in the genome, are not used. All the reads parsing were done with samtools. **Figure S1** shows the flow chart for the filtering, mapping, and identification of PCH Hi-C reads, and the number of reads at each step. Genome-wide contact maps for both PCH and euchromatic regions (**Figure S4 and S5**) were generated using HOMER with simple normalization [94].

#### Spatial interaction between PCH regions

Hi-C read pairs whose both ends mapped uniquely to epigenetically defined PCH were included in the analysis. Read pairs whose mapping locations are within 10kb to each other were removed, as our analysis focuses on long-range spatial interactions. We performed three sequential analyses (all read pairs, excluding intra-arm read pairs, excluding intra-chromosome read pairs) to identify three types of PCH-PCH interactions: within arm, within chromosome between arms (e.g. 2L-2R, 3L-3R), and between chromosomes. It is worth noting that not enough sequences have been assembled on the short arms of X, Y and 4^th^ chromosomes, thus precluding within chromosome, between arms analysis for these chromosomes. Theoretical percentage of each pairwise interaction among PCH regions on different chromosomes was estimated based on the mappability track of *D. melanogaster* Release 6 genome, which was generated using GEM mappability tool (using read length 50 and other default parameters, [40]). We then counted the number of bases with mappability one (i.e. can be unambiguously mapped in the genome) in the PCH regions of each chromosome. Empirical expected percentage of each pairwise interaction was estimated from the percentage of reads mapping uniquely to the PCH on each chromosome arm, ignoring read pair information. Because the Hi-C data were generated using unsexed embryos, we assumed equal sex ratio when estimating expectations. To assess whether the observed percentage is more than the empirical expectation, we randomly permuted 10,000 times read pair labels, generated an empirical distribution of the percentage, and calculated one-sided *p-values*.

#### Spatial interaction between euchromatic regions and heterochromatin

We used samtools to parse out read pairs whose one end mapped uniquely (with mapping quality at least 30) within the focused euchromatin regions, and estimated the percentage of PCH reads at the other end. All three categories of heterochromatic reads were included. Regions with less than 1,000 Hi-C read pairs were excluded from the analysis. We found strong correlations between replicates for both the percentage of euchromatin-PCH reads and the associated *p-values* (see below) for H3K9me2-enriched regions and TEs (*Spearman rank ρ* > 88%, *p* < 10^-16^, **Figure S22 and S23**). To assess whether the percentage of euchromatin-PCH read pairs is significant, we randomly selected euchromatic regions without H3K9me2 enrichment, performed the same analysis to get a null distribution of the percentage, and estimated the *p-values.* We simulated 200 sets of non-H3K9me2 enriched random euchromatic regions that are of the same sample size, on the same chromosome and, for H3K9me2 enriched regions, of the same size as the focused set. This was done separately for H3K9me2 enriched regions and TEs, and separately for the two replicates. Because of the tendency of within chromosome interactions (see main text) and the difference in the PCH sizes among chromosomes [36, 37], the percentage of euchromatin-PCH read pairs varies between randomly selected regions on different chromosomes (**Figure S24**). Also, smaller euchromatic regions have fewer Hi-C read pairs included in the analysis, which translates into smaller sample size and thus larger variance of the estimated percentage (**Figure S25**), leading to the estimates more likely to hit the boundary condition (i.e. no euchromatin-PCH read pairs, **Figure S25**, red circles). Accordingly, for each euchromatic region, the *p-value* is estimated using random regions that are on the same chromosome and of the same size quantile. For H3K9me2-enriched euchromatic regions, we used +/-1kb of the enriched region as the defined window. Because, unlike H3K9me2 enriched regions, euchromatic TEs were identified as a small interval with possible insertions within [83], we used +/− 2kb of the TE insertion site/interval as the defined window.

### Generation of FISH probes

#### Heterochromatic repeat probes

LNA probes [95] targeting AAGAG (bulk heterochromatin), AACAC (2R PCH), dodeca (3R PCH), AATAT (4^th^ and Y), and AATAGAC (Y) were ordered from Integrated DNA Technologies (IDT).

#### Oligopaint FISH probes

We designed Oligopaint probes that target single copy genome regions, following [44, 96]. Each targeted euchromatic region has at least 500 probes designed to label it, with at least 12 probes/kb (**Table S5**). For euchromatic TEs, designed Oligopaint probes target the “flanking” unique sequences instead of the TE itself. Within the total oligo library, each pool of probes targeting a genomic region was designed with an appended specific barcode (https://github.com/gnir/OligoLego, [97]), and an additional Universal barcodes that were appended to the very 5’ and 3’ ends, both were used for PCR amplification of the specific/total library. Primary Oligopaint libraries were ordered from CustomArray (Bothell, WA), and amplified and synthesized into Oligopaint probes following [46]. To label specific subsets of oligos within the library, complementary “bridge” oligos were hybridized against their barcodes, followed by hybridization with fluorophore-labeled secondary oligos complementary to an overhang of the bridge oligo. Bridge oligos and fluorescence labeled secondary Oligopaint probe were ordered from IDT.

### Embryo collections, treatments, and fixations

#### Embryo collections

Flies laid eggs on fresh apple juice plate for 1hr (pre-lay), followed by 2hr egg-laying on new apple juice plates. Collected embryos were incubated at 25°C for 16hr to harvest 16-18hr embryos, which were then fixed immediately.

#### Embryo permealization and 1,6-hexanediol treatment

To allow effective permeabilization of 16-18hr late stage embryos for 1,6-hexanediol treatment, 0-2hr embryos were incubated at 18°C for 32hr, which equals to 16hr development at 25°C [98]. Embryos were dechorionated in 50% bleach for 90s, washed with water for 1 min, and treated with EPS, a d-limonene based solvent with low toxicity [98, 99], for 2 min. Permeabilized embryos were either fixed immediately or incubated in 10% 1,6-hexanediol (dissolved in PBS) for 4 min, followed by a quick wash with PBS and fixed immediately.

#### Fixation of embryos

16-18hr embryos (without treatment) were dechorionated in 50% bleach for 90s, washed with water for 1 min. Dechorionated embryos, embryos with EPS treatment, or embryos with EPS and 1,6-hexanediol treatments were transferred to biphasic fixation solution with 4% formaldehyde (1.2mL Heptane, 75μL 16% formaldehyde, and 225 μL PBS), and shake for 20 min at room temperature. Embryos were then transferred to tubes with biphasic solution of equal volume of heptane and methanol, followed by vigorous shaking for 30-45s to crack the embryos, three washed with methanol, and stored in −20°C in methanol.

### FISH

#### Repeat probes

Embryos (stored in methanol) were rehydrated sequentially into PBT (1xPBS, 0.1% Tween-20), incubated with 100 μg/mL RNAseA in PBT for two hours at room temperature, washed twice with PBT, post-fixed with 4% formaldehyde in PBT for 20min, washed three times with PBT, and then sequentially transitioned into hybridization buffer (50% formamide, 5x SSC, 100 μg/ml Heparin, 100 μg/ml sheared salmon sperm DNA, and 0.1% Tween-20). Before hybridization, embryos were incubated with pre-hybridization solution (hybridization buffer boiled at 100°C for 5 min, chilled on ice) at 56°C for at least two hours. Embryos were then incubated with 25 ng/μL of LNA repeat probes (denatured at 70°C for 3 min) at 80°C for 15 min and then 37°C with shaking overnight. For FISH with AATAT probe, embryos were incubated at 37°C for three hours, then 25°C overnight. Embryos were washed with hybridization buffer twice at 37/25°C, followed by sequential transition into PBT, two PBT washes at room temperature, DAPI staining, two PBS washes, resuspended in Prolong Gold Antifade (Life Technologies), and mounted on slides.

We used AATAT to mark 4^th^ chromosome heterochromatin. Because this repeat is also abundant on the Y [41], embryos were also stained with Y-specific repeat, AATAGAC and female embryos were analyzed.

#### Oligopaint probes and AAGAG probe

Embryo FISH with both Oligopaint and AAGAG (for bulk heterochromatin) LNA probe followed [100], except for staining nuclei with DAPI and resuspension in Prolong Gold Antifade (Life Technologies).

### Imaging and data analysis

Images of embryos were collected on Zeiss LSM710 confocal florescence microscope, using a 1.4NA 63X oil objective (Zeiss), and analyzed in Fiji [101]. Distances between foci were measured by Fiji linetool, and divided by the radius of the nucleus to get relative distance. In cases where the nuclei are not perfectly round, we used radius on the longest axis. There are usually one to two AAGAG foci in a nucleus and the distance was measured between Oligopaint focus and the nearest AAGAG focus. At least 70 nuclei were counted for each treatment/genotype.

## Data availability

Genome sequence (raw genome data and called transposable elements) and ChIP-seq data (raw data and processed tracks) have been deposited to GEO (GSE125307 – private token for reviewers: gzavmcqizxmjxil; GSE125031 – private token for reviewers: mrmhquucztcdlwv).

## Acknowledgements

We thank Kyle Millis for technical help with FISH, members of the Karpen lab for many helpful discussions, and Aniek Janssen, Serafin Colmenares, and Sasha Langley for carefully reading the manuscript. We thank Charles Langley for providing computational resources. We appreciate Jumana AlHaj Abed and Jeleca Erceg for helpful discussions of the Oligopaint experimental design. We also thank Vincent J. Coates Genomics Sequencing Laboratory (GSL) at UC Berkeley, MGX sequencing and Drosophila facilities (BioCampus Montpellier, CNRS, INSERM, Université de Montpellier).

## Author contributions

Conceptualization – YCGL, GHK; Data curation – YCGL, YO; Formal analysis – YCGL; Funding acquisition – CTW, GC, GHK; Investigation – YCGL, YO, DA; Methodology – YCGL, NMCM, BJB; Resources – CTW, GC, GHK; Supervision – GHK; Writing – original draft – YCGL; Writing – review & editing – YCGL, YO, NMCM, BJB, CTW, GC, GHK.

## Fudning

YCGL is supported by NIH K99 GM121868, YO is supported by JSPS Overseas Research Fellowships (Japan Society for the Promotion of Science), BJB is supported by Damon Runyon Cancer Research Fellowship (HHMI), CTW is supported by NIH DP1GM106412, RO1HD091797, and RO1GM123289, GC is supported by European Research Council (ERC-2008-AdG no. 232947), the CNRS, the European Union’s Horizon 2020 research and innovation program under grant agreement No. 676556 (MuG), and the Agence Nationale de la Recherche (N. ANR-15-CE12-006-01), and GHK is supported by NIH R01 GM117420. All authors declare no competing interests.

## Supplementary files

**Figure S1. Flow chart for identification of PCH Hi-C reads**

**Figure S2. Heatmap for the number of Hi-C read pairs supporting the spatial interactions between pairs of 100kb PCH windows for Hi-C replicate 2.** Note that only the PCH regions are shown.

**Figure S3. Circular plot showing inter-arm and inter-chromosomal interactions.** Circular plot showing inter-arm and inter-chromosome interactions supported by 95, 99, and 99.9 percentile of Hi-C reads. Average mappability of each window is shown in the inner track.

**Figure S4. Genome-wide normalized contact map of replicate 1.** Both unique euchromatic and PCH regions are shown. Blue bars are PCH regions while gray bars are euchromatic regions. Centromeres are denoted as triangles. Each element in the matrix represents the *log ratio* between the number of observed contact (Hi-C read pairs) and the number expected under the assumption that each 500kb window would have equal number of total interactions across the genome. The number of observed contacts involving Y chromosome is too low for proper normalization and thus excluded from representation in the figure. Note that this normalization may be biased against interactions involving PCH regions (EU-PCH and PCH-PCH) because much fewer reads uniquely mapped to PCH regions than euchromatic regions.

**Figure S5. Genome-wide normalized contact map of replicate 2.**

**Figure S6. FISH validation for identified PCH-PCH interaction using absolute distance.** (A) Boxplot and (B) histograms showing the absolute distance between PCH foci. Comparisons of distance between pairs of foci were tested with *Mann-Whitney test* (*p-values* in (A)) and *Fisher’s exact test* (for proportion of overlapping foci, *p-values* = 0.001 (3R-4 vs 2R-4) and 0.22 (3R-4 vs 3R-2R)). Threshold for nuclei with overlapping foci is denoted with arrow, which corresponds to 0.1 μm.

**Figure S7. Boxplot of the linear distance to PCH for H3K9me2 islands.** H3K9me2-enriched with and without PCH interactions are in green and gray respectively.

**Figure S8. Percentage of uniquely mapped heterochromatic Hi-C reads coming from a particular chromosome for euchromatic regions on different chromosomes.** Data for replicate 2 is shown.

**Figure S9. H3K9me2 enrichment level for euchromatic regions chosen for FISH analysis.** There is H3K9me2 enrichment in both ORw1118 and wildtype strains for EU1-3, but none for control regions c.EU1-3. The fourth tracks (below RAL360, blue) are broad peaks called by Macs2 in ORw1118.

**Figure S10. Representative FISH images for chosen euchromatic regions and PCH.**

**Figure S11. FISH validation for identified EU-PCH interaction using absolute distance.** (A) Boxplot and (B) histogram showing the absolute distance between euchromatic loci and PCH. Comparisons of distance between pairs of foci were tested with *Mann-Whitney test* (*p-values* in (A)) and *Fisher’s exact test* (for proportion of overlapping foci, *p-values* = 0.01 (EU1 vs c.EU1), 0.53 (EU2 vs c.EU2), and 0.15 (EU3 vs c.EU3)). Threshold for nuclei with overlapping foci is denoted with arrow, which corresponds to 0.1 μm.

**Figure S12. H3K9me2 enrichment level in euchromatic TE neighborhood.** Strain-specific H3K9me2 enrichment was observed for TE1 and TE2. Third track (one below RAL315, green) shows the insertion position of TEs in ORw1118 predicted by TIDAL.

**Figure S13. Genomic distribution of TEs with and without PCH interactions.** The extent of local H3K9me2 enrichment at TEs is shown on the y-axis for TEs with (green) and without (gray) local H3K9me2 enrichment, and with (dark green) and without (light green) PCH interaction.

**Figure S14. Boxplot of the linear distance to PCH for euchromatic TEs.** TEs with and without PCH interactions are in dark and light green respectively.

**Figure S15. FISH validation for identified TE-PCH interaction using absolute distance.** (A) Boxplot and (B) histogram showing the absolute distance between euchromatic TEs and PCH. Comparisons of distance between pairs of foci were tested with *Mann-Whitney test* (*p-values* in (A)) and *Fisher’s exact test* (for proportion of overlapping foci, *p-values* = 0.0007 (TE1), 0.07 (TE2), 0.48 (c.TE1), and 1 (c.TE2)). Threshold for nuclei with overlapping foci is denoted with arrow, which corresponds to 0.1 μm.

**Figure S16. Representative FISH images for permealized embryos (EPS) and permealized embryos with 1,6-hexanediol treatment (EPS+HD).**

**Figure S17. FISH validation for the influence of 1,6-Hexanediol on the spatial associations between euchromatic TE and PCH using absolute distance.** (A) Boxplot and (B) histogram showing the absolute distance between euchromatic TE and PCH. Comparisons of distance between pairs of foci were tested with *Mann-Whitney test* (*p-values* in (A)) and *Fisher’s exact test* (for proportion of overlapping foci, *p-values* = 0.02 (ORw, EPS vs EPS+HD), 1 (WT, EPS vs EPS+HD), 0.057 (ESP treatment, ORw vs WT), 0.55 (HD treatment, ORw vs WT)). Threshold for nuclei with overlapping foci is denoted with arrow, which corresponds to 0.1 μm.

**Figure S18. Extent and magnitude of H3K9me2 enrichment of TEs with and without PCH interaction.**

**Figure S19. IDR plots for ORw1118**

**Figure S20. IDR plots for RAL315**

**Figure S21. IDR plots for RAL360**

**Figure S22. X-Y plots for the estimated proportion of euchromatin-PCH reads, and the associated *p-values*, between Hi-C replicates for TEs.**

**Figure S23. X-Y plots for the estimated proportion of euchromatin-PCH reads, and the associated *p-values*, between Hi-C replicates for H3K9me2-enriched regions.**

**Figure S24. Distribution of estimated euchromatin-PCH read pairs for random regions on different chromosomes.**

**Figure S25. Distribution of estimated euchromatin-PCH read pairs for random regions of different size.**

**Table S1.** List of heterochromatic simple repeats

**Table S2.** Chromatin environment of euchromatic H3K9me2-enriched regions interacting with PCH

**Table S3.** Properties of euchromatic H3K9me2-enriched regions interacting with PCH

**Table S4.** Information for regions targeted by Oligopaint

**Table S5.** Properties of euchromatic TEs interacting with PCH

**Table S6.** Trans epigenetic effects of TEs

## References

1. Misteli T, Soutoglou E. The emerging role of nuclear architecture in DNA repair and genome maintenance. Nat Rev Mol Cell Biol. 2009;10: 243–254. doi:10.1038/nrm2651

2. Bickmore WA, van Steensel B. Genome Architecture: Domain Organization of Interphase Chromosomes. Cell. 2013;152: 1270–1284. doi:10.1016/j.cell.2013.02.001

3. Sexton T, Cavalli G. The Role of Chromosome Domains in Shaping the Functional Genome. Cell. 2015;160: 1049–1059. doi:10.1016/j.cell.2015.02.040

4. Bonev B, Cavalli G. Organization and function of the 3D genome. Nat Rev Genet. 2016;17: 661–678. doi:10.1038/nrg.2016.112

5. Dekker J, Rippe K, Dekker M, Kleckner N. Capturing Chromosome Conformation. Science. 2002;295: 1306–1311. doi:10.1126/science.1067799

6. Lieberman-Aiden E, Berkum NL van, Williams L, Imakaev M, Ragoczy T, Telling A, et al. Comprehensive Mapping of Long-Range Interactions Reveals Folding Principles of the Human Genome. Science. 2009;326: 289–293. doi:10.1126/science.1181369

7. Denker A, Laat W de. The second decade of 3C technologies: detailed insights into nuclear organization. Genes Dev. 2016;30: 1357–1382. doi:10.1101/gad.281964.116

8. Sexton T, Yaffe E, Kenigsberg E, Bantignies F, Leblanc B, Hoichman M, et al. Three-Dimensional Folding and Functional Organization Principles of the Drosophila Genome. Cell. 2012;148: 458–472. doi:10.1016/j.cell.2012.01.010

9. Dixon JR, Selvaraj S, Yue F, Kim A, Li Y, Shen Y, et al. Topological domains in mammalian genomes identified by analysis of chromatin interactions. Nature. 2012 [cited 11 Apr 2012]. doi:10.1038/nature11082

10. Stadler MR, Haines JE, Eisen MB. Convergence of topological domain boundaries, insulators, and polytene interbands revealed by high-resolution mapping of chromatin contacts in the early Drosophila melanogaster embryo. In: eLife [Internet]. 17 Nov 2017 [cited 7 Sep 2018]. Available: https://elifesciences.org/articles/29550

11. Arabidopsis Genome Initiative. Analysis of the genome sequence of the flowering plant Arabidopsis thaliana. Nature. 2000;408: 796–815. doi:10.1038/35048692

12. Smith CD, Shu S, Mungall CJ, Karpen GH. The Release 5.1 Annotation of Drosophila melanogaster Heterochromatin. Science. 2007;316: 1586 –1591. doi:10.1126/science.1139815

13. Karpen GH, Le M-H, Le H. Centric Heterochromatin and the Efficiency of Achiasmate Disjunction in Drosophila Female Meiosis. Science. 1996;273: 118–122. doi:10.1126/science.273.5271.118

14. Dernburg AF, Sedat JW, Hawley RS. Direct evidence of a role for heterochromatin in meiotic chromosome segregation. Cell. 1996;86: 135–146.

15. Peters AHFM, O’Carroll D, Scherthan H, Mechtler K, Sauer S, Schöfer C, et al. Loss of the Suv39h Histone Methyltransferases Impairs Mammalian Heterochromatin and Genome Stability. Cell. 2001;107: 323–337. doi:10.1016/S0092-8674(01)00542-6

16. Peng JC, Karpen GH. Heterochromatic Genome Stability Requires Regulators of Histone H3 K9 Methylation. PLOS Genet. 2009;5: e1000435. doi:10.1371/journal.pgen.1000435

17. Janssen A, Colmenares SU, Karpen GH. Heterochromatin: Guardian of the Genome. Annu Rev Cell Dev Biol. 2018;34: 265–288. doi:10.1146/annurev-cellbio-100617-062653

18. James TC, Elgin SC. Identification of a nonhistone chromosomal protein associated with heterochromatin in Drosophila melanogaster and its gene. Mol Cell Biol. 1986;6: 3862– 3872. doi:10.1128/MCB.6.11.3862

19. Jacobs SA, Taverna SD, Zhang Y, Briggs SD, Li J, Eissenberg JC, et al. Specificity of the HP1 chromo domain for the methylated N-terminus of histone H3. EMBO J. 2001;20: 5232– 5241. doi:10.1093/emboj/20.18.5232

20. Zhang P, Spradling AC. The Drosophila Salivary Gland Chromocenter Contains Highly Polytenized Subdomains of Mitotic Heterochromatin. Genetics. 1995;139: 659–670.

21. Mayer R, Brero A, von Hase J, Schroeder T, Cremer T, Dietzel S. Common themes and cell type specific variations of higher order chromatin arrangements in the mouse. BMC Cell Biol. 2005;6: 44. doi:10.1186/1471-2121-6-44

22. Larson AG, Elnatan D, Keenen MM, Trnka MJ, Johnston JB, Burlingame AL, et al. Liquid droplet formation by HP1α suggests a role for phase separation in heterochromatin. Nature. 2017;547: 236–240. doi:10.1038/nature22822

23. Strom AR, Emelyanov AV, Mir M, Fyodorov DV, Darzacq X, Karpen GH. Phase separation drives heterochromatin domain formation. Nature. 2017;547: 241–245. doi:10.1038/nature22989

24. Wakimoto BT, Hearn MG. The effects of chromosome rearrangements on the expression of heterochromatic genes in chromosome 2L of Drosophila melanogaster. Genetics. 1990;125: 141–154.

25. Gowen JW, Gay EH. EFFECT OF TEMPERATURE ON EVERSPORTING EYE COLOR IN DROSOPHILA MELANOGASTER. Science. 1933;77: 312. doi:10.1126/science.77.1995.312

26. Girton JR, Johansen KM. Chromatin structure and the regulation of gene expression: the lessons of PEV in Drosophila. Adv Genet. 2008;61: 1–43. doi:10.1016/S0065-2660(07)00001-6

27. Elgin SCR, Reuter G. Position-effect variegation, heterochromatin formation, and gene silencing in Drosophila. Cold Spring Harb Perspect Biol. 2013;5: a017780. doi:10.1101/cshperspect.a017780

28. Falk M, Feodorova Y, Naumova N, Imakaev M, Lajoie BR, Leonhardt H, et al. Heterochromatin drives compartmentalization of inverted and conventional nuclei. Nature. 2019;570: 395. doi:10.1038/s41586-019-1275-3

29. Vogel MJ, Guelen L, de Wit E, Hupkes DP, Lodén M, Talhout W, et al. Human heterochromatin proteins form large domains containing KRAB-ZNF genes. Genome Res. 2006;16: 1493–1504. doi:10.1101/gr.5391806

30. Wen B, Wu H, Shinkai Y, Irizarry RA, Feinberg AP. Large histone H3 lysine 9 dimethylated chromatin blocks distinguish differentiated from embryonic stem cells. Nat Genet. 2009;41: 246–250. doi:10.1038/ng.297

31. Riddle NC, Minoda A, Kharchenko PV, Alekseyenko AA, Schwartz YB, Tolstorukov MY, et al. Plasticity in patterns of histone modifications and chromosomal proteins in Drosophila heterochromatin. Genome Res. 2011;21: 147–163. doi:10.1101/gr.110098.110

32. Dernburg AF, Broman KW, Fung JC, Marshall WF, Philips J, Agard DA, et al. Perturbation of Nuclear Architecture by Long-Distance Chromosome Interactions. Cell. 1996;85: 745– 759. doi:10.1016/S0092-8674(00)81240-4

33. Csink AK, Henikoff S. Genetic modification of heterochromatic association and nuclear organization in Drosophila. Nature. 1996;381: 529–531. doi:10.1038/381529a0

34. Lee YCG. The Role of piRNA-Mediated Epigenetic Silencing in the Population Dynamics of Transposable Elements in Drosophila melanogaster. PLoS Genet. 2015;11: e1005269. doi:10.1371/journal.pgen.1005269

35. Lee YCG, Karpen GH. Pervasive epigenetic effects of Drosophila euchromatic transposable elements impact their evolution. eLife. 2017;6. doi:10.7554/eLife.25762

36. Hoskins RA, Carlson JW, Kennedy C, Acevedo D, Evans-Holm M, Frise E, et al. Sequence Finishing and Mapping of Drosophila melanogaster Heterochromatin. Science. 2007;316: 1625 –1628. doi:10.1126/science.1139816

37. Hoskins RA, Carlson JW, Wan KH, Park S, Mendez I, Galle SE, et al. The Release 6 reference sequence of the Drosophila melanogaster genome. Genome Res. 2015; gr.185579.114. doi:10.1101/gr.185579.114

38. Schuettengruber B, Oded Elkayam N, Sexton T, Entrevan M, Stern S, Thomas A, et al. Cooperativity, specificity, and evolutionary stability of Polycomb targeting in Drosophila. Cell Rep. 2014;9: 219–233. doi:10.1016/j.celrep.2014.08.072

39. Gatti M, Pimpinelli S. Functional Elements in Drosophila Melanogaster Heterochromatin. Annu Rev Genet. 1992;26: 239–276. doi:10.1146/annurev.ge.26.120192.001323

40. Derrien T, Estellé J, Sola SM, Knowles DG, Raineri E, Guigó R, et al. Fast Computation and Applications of Genome Mappability. PLOS ONE. 2012;7: e30377. doi:10.1371/journal.pone.0030377

41. Dernburg AF. In Situ Hybridization to Somatic Chromosomes in Drosophila. Cold Spring Harb Protoc. 2011;2011: pdb.top065540. doi:10.1101/pdb.top065540

42. Filion GJ, van Bemmel JG, Braunschweig U, Talhout W, Kind J, Ward LD, et al. Systematic protein location mapping reveals five principal chromatin types in Drosophila cells. Cell. 2010;143: 212–224. doi:10.1016/j.cell.2010.09.009

43. Kharchenko PV, Alekseyenko AA, Schwartz YB, Minoda A, Riddle NC, Ernst J, et al. Comprehensive analysis of the chromatin landscape in Drosophila melanogaster. Nature. 2011;471: 480–485. doi:10.1038/nature09725

44. Beliveau BJ, Joyce EF, Apostolopoulos N, Yilmaz F, Fonseka CY, McCole RB, et al. Versatile design and synthesis platform for visualizing genomes with Oligopaint FISH probes. Proc Natl Acad Sci U S A. 2012;109: 21301–21306. doi:10.1073/pnas.1213818110

45. Beliveau BJ, Apostolopoulos N, Wu C. Visualizing genomes with Oligopaint FISH probes. Curr Protoc Mol Biol. 2014;105: Unit 14.23. doi:10.1002/0471142727.mb1423s105

46. Beliveau BJ, Boettiger AN, Nir G, Bintu B, Yin P, Zhuang X, et al. In Situ Super-Resolution Imaging of Genomic DNA with OligoSTORM and OligoDNA-PAINT. In: Erfle H, editor. Super-Resolution Microscopy: Methods and Protocols. New York, NY: Springer New York; 2017. pp. 231–252. Available: https://doi.org/10.1007/978-1-4939-7265-4_19

47. Lohe AR, Brutlag DL. Multiplicity of satellite DNA sequences in Drosophila melanogaster. Proc Natl Acad Sci U S A. 1986;83: 696–700.

48. Lohe AR, Hilliker AJ, Roberts PA. Mapping Simple Repeated DNA Sequences in Heterochromatin of Drosophila Melanogaster. Genetics. 1993;134: 1149–1174.

49. Rebollo R, Karimi MM, Bilenky M, Gagnier L, Miceli-Royer K, Zhang Y, et al. Retrotransposon-Induced Heterochromatin Spreading in the Mouse Revealed by Insertional Polymorphisms. PLoS Genet. 2011;7: e1002301. doi:10.1371/journal.pgen.1002301

50. Sentmanat MF, Elgin SCR. Ectopic assembly of heterochromatin in Drosophila melanogaster triggered by transposable elements. Proc Natl Acad Sci. 2012;109: 14104–14109. doi:10.1073/pnas.1207036109

51. Charlesworth B, Langley CH. The population genetics of Drosophila transposable elements. Annu Rev Genet. 1989;23: 251–287. doi:10.1146/annurev.ge.23.120189.001343

52. Cridland JM, Macdonald SJ, Long AD, Thornton KR. Abundance and Distribution of Transposable Elements in Two Drosophila QTL Mapping Resources. Mol Biol Evol. 2013;30: 2311–2327. doi:10.1093/molbev/mst129

53. Kofler R, Nolte V, Schlötterer C. Tempo and Mode of Transposable Element Activity in Drosophila. PLoS Genet. 2015;11: e1005406. doi:10.1371/journal.pgen.1005406

54. Park PJ. ChIP–seq: advantages and challenges of a maturing technology. Nat Rev Genet. 2009;10: 669–680. doi:10.1038/nrg2641

55. Nakato R, Shirahige K. Recent advances in ChIP-seq analysis: from quality management to whole-genome annotation. Brief Bioinform. 2017;18: 279–290. doi:10.1093/bib/bbw023

56. Ribbeck K, Görlich D. The permeability barrier of nuclear pore complexes appears to operate via hydrophobic exclusion. EMBO J. 2002;21: 2664–2671. doi:10.1093/emboj/21.11.2664

57. Lee YCG, Langley CH. Transposable elements in natural populations of Drosophila melanogaster. Philos Trans R Soc B Biol Sci. 2010;365: 1219–1228. doi:10.1098/rstb.2009.0318

58. Barrón MG, Fiston-Lavier A-S, Petrov DA, González J. Population Genomics of Transposable Elements in Drosophila. Annu Rev Genet. 2014;48: 561–581. doi:10.1146/annurev-genet-120213-092359

59. Lack JB, Cardeno CM, Crepeau MW, Taylor W, Corbett-Detig RB, Stevens KA, et al. The Drosophila Genome Nexus: A Population Genomic Resource of 623 Drosophila melanogaster Genomes, Including 197 from a Single Ancestral Range Population. Genetics. 2015; genetics.115.174664. doi:10.1534/genetics.115.174664

60. Petrov DA, Fiston-Lavier A-S, Lipatov M, Lenkov K, González J. Population genomics of transposable elements in Drosophila melanogaster. Mol Biol Evol. 2011;28: 1633–1644. doi:10.1093/molbev/msq337

61. Lander ES, Linton LM, Birren B, Nusbaum C, Zody MC, Baldwin J, et al. Initial sequencing and analysis of the human genome. Nature. 2001;409: 860–921. doi:10.1038/35057062

62. Lichter P, Cremer T, Borden J, Manuelidis L, Ward DC. Delineation of individual human chromosomes in metaphase and interphase cells by in situ suppression hybridization using recombinant DNA libraries. Hum Genet. 1988;80: 224–234. doi:10.1007/BF01790090

63. Pinkel D, Landegent J, Collins C, Fuscoe J, Segraves R, Lucas J, et al. Fluorescence in situ hybridization with human chromosome-specific libraries: detection of trisomy 21 and translocations of chromosome 4. Proc Natl Acad Sci U S A. 1988;85: 9138–9142.

64. Kalhor R, Tjong H, Jayathilaka N, Alber F, Chen L. Genome architectures revealed by tethered chromosome conformation capture and population-based modeling. Nat Biotechnol. 2012;30: 90–98. doi:10.1038/nbt.2057

65. Williamson I, Berlivet S, Eskeland R, Boyle S, Illingworth RS, Paquette D, et al. Spatial genome organization: contrasting views from chromosome conformation capture and fluorescence in situ hybridization. Genes Dev. 2014;28: 2778–2791. doi:10.1101/gad.251694.114

66. Rosin LF, Nguyen SC, Joyce EF. Condensin II drives large-scale folding and spatial partitioning of interphase chromosomes in Drosophila nuclei. PLOS Genet. 2018;14: e1007393. doi:10.1371/journal.pgen.1007393

67. Dekker J, Marti-Renom MA, Mirny LA. Exploring the three-dimensional organization of genomes: interpreting chromatin interaction data. Nat Rev Genet. 2013;14: 390–403. doi:10.1038/nrg3454

68. Haddad N, Jost D, Vaillant C. Perspectives: using polymer modeling to understand the formation and function of nuclear compartments. Chromosome Res. 2017;25: 35–50. doi:10.1007/s10577-016-9548-2

69. Swenson JM, Colmenares SU, Strom AR, Costes SV, Karpen GH. The composition and organization of Drosophila heterochromatin are heterogeneous and dynamic. eLife. 2016;5: e16096. doi:10.7554/eLife.16096

70. Hearn MG, Hedrick A, Grigliatti TA, Wakimoto BT. The effect of modifiers of position-effect variegation on the variegation of heterochromatic genes of Drosophila melanogaster. Genetics. 1991;128: 785–797.

71. Yasuhara JC, Wakimoto BT. Molecular Landscape of Modified Histones in Drosophila Heterochromatic Genes and Euchromatin-Heterochromatin Transition Zones. PLoS Genet. 2008;4: e16. doi:10.1371/journal.pgen.0040016

72. Piacentini L, Fanti L, Berloco M, Perrini B, Pimpinelli S. Heterochromatin protein 1 (HP1) is associated with induced gene expression in Drosophila euchromatin. J Cell Biol. 2003;161: 707–714. doi:10.1083/jcb.200303012

73. Piacentini L, Fanti L, Negri R, Vescovo VD, Fatica A, Altieri F, et al. Heterochromatin Protein 1 (HP1a) Positively Regulates Euchromatic Gene Expression through RNA Transcript Association and Interaction with hnRNPs in Drosophila. PLOS Genet. 2009;5: e1000670. doi:10.1371/journal.pgen.1000670

74. Cryderman DE, Grade SK, Li Y, Fanti L, Pimpinelli S, Wallrath LL. Role of Drosophila HP1 in euchromatic gene expression. Dev Dyn. 2005;232: 767–774. doi:10.1002/dvdy.20310

75. Joyce EF, Erceg J, Wu C -tin. Pairing and anti-pairing: a balancing act in the diploid genome. Curr Opin Genet Dev. 2016;37: 119–128. doi:10.1016/j.gde.2016.03.002

76. Henikoff S, Dreesen TD. Trans-inactivation of the Drosophila brown gene: evidence for transcriptional repression and somatic pairing dependence. Proc Natl Acad Sci. 1989;86: 6704–6708.

77. Elliott TA, Gregory TR. Do larger genomes contain more diverse transposable elements? BMC Evol Biol. 2015;15: 69. doi:10.1186/s12862-015-0339-8

78. Consortium IHGS. Finishing the euchromatic sequence of the human genome. Nature. 2004;431: 931–945. doi:10.1038/nature03001

79. Stewart C, Kural D, Strömberg MP, Walker JA, Konkel MK, Stütz AM, et al. A Comprehensive Map of Mobile Element Insertion Polymorphisms in Humans. PLoS Genet. 2011;7: e1002236. doi:10.1371/journal.pgen.1002236

80. Sudmant PH, Rausch T, Gardner EJ, Handsaker RE, Abyzov A, Huddleston J, et al. An integrated map of structural variation in 2,504 human genomes. Nature. 2015;526: 75– 81. doi:10.1038/nature15394

81. Dolgin ES, Charlesworth B, Cutter AD. Population frequencies of transposable elements in selfing and outcrossing Caenorhabditis nematodes. Genet Res. 2008;90: 317–329. doi:10.1017/S0016672308009440

82. Laricchia KM, Zdraljevic S, Cook DE, Andersen EC. Natural Variation in the Distribution and Abundance of Transposable Elements Across the Caenorhabditis elegans Species. Mol Biol Evol. 2017;34: 2187–2202. doi:10.1093/molbev/msx155

83. Rahman R, Chirn G, Kanodia A, Sytnikova YA, Brembs B, Bergman CM, et al. Unique transposon landscapes are pervasive across Drosophila melanogaster genomes. Nucleic Acids Res. 2015;43: 10655–10672. doi:10.1093/nar/gkv1193

84. Wright SI, Le QH, Schoen DJ, Bureau TE. Population dynamics of an Ac-like transposable element in self- and cross-pollinating arabidopsis. Genetics. 2001;158: 1279–1288.

85. Quadrana L, Silveira AB, Mayhew GF, LeBlanc C, Martienssen RA, Jeddeloh JA, et al. The Arabidopsis thaliana mobilome and its impact at the species level. eLife. 2016;5: e15716. doi:10.7554/eLife.15716

86. Stuart T, Eichten SR, Cahn J, Karpievitch YV, Borevitz JO, Lister R. Population scale mapping of transposable element diversity reveals links to gene regulation and epigenomic variation. eLife. 2016;5: e20777. doi:10.7554/eLife.20777

87. Mackay TFC, Richards S, Stone EA, Barbadilla A, Ayroles JF, Zhu D, et al. The Drosophila melanogaster Genetic Reference Panel. Nature. 2012;482: 173–178. doi:10.1038/nature10811

88. Babraham Bioinformatics - Trim Galore! [cited 18 Nov 2016]. Available: http://www.bioinformatics.babraham.ac.uk/projects/trim_galore/

89. Li H. A statistical framework for SNP calling, mutation discovery, association mapping and population genetical parameter estimation from sequencing data. Bioinforma Oxf Engl. 2011;27: 2987–2993. doi:10.1093/bioinformatics/btr509

90. Zhang Y, Liu T, Meyer CA, Eeckhoute J, Johnson DS, Bernstein BE, et al. Model-based Analysis of ChIP-Seq (MACS). Genome Biol. 2008;9: R137. doi:10.1186/gb-2008-9-9-r137

91. Li Q, Brown JB, Huang H, Bickel PJ. Measuring reproducibility of high-throughput experiments. Ann Appl Stat. 2011;5: 1752–1779. doi:10.1214/11-AOAS466

92. Kaminker JS, Bergman CM, Kronmiller B, Carlson J, Svirskas R, Patel S, et al. The transposable elements of the Drosophila melanogaster euchromatin: a genomics perspective. Genome Biol. 2002;3: RESEARCH0084.

93. Li H, Durbin R. Fast and accurate long-read alignment with Burrows-Wheeler transform. Bioinformatics. 2010;26: 589–595. doi:10.1093/bioinformatics/btp698

94. Heinz S, Benner C, Spann N, Bertolino E, Lin YC, Laslo P, et al. Simple combinations of lineage-determining transcription factors prime cis-regulatory elements required for macrophage and B cell identities. Mol Cell. 2010;38: 576–589. doi:10.1016/j.molcel.2010.05.004

95. Williams BR, Bateman JR, Novikov ND, Wu C-T. Disruption of Topoisomerase II Perturbs Pairing in Drosophila Cell Culture. Genetics. 2007;177: 31–46. doi:10.1534/genetics.107.076356

96. Beliveau BJ, Kishi JY, Nir G, Sasaki HM, Saka SK, Nguyen SC, et al. OligoMiner provides a rapid, flexible environment for the design of genome-scale oligonucleotide in situ hybridization probes. Proc Natl Acad Sci U S A. 2018;115: E2183–E2192. doi:10.1073/pnas.1714530115

97. Nir G, Farabella I, Estrada CP, Ebeling CG, Beliveau BJ, Sasaki HM, et al. Walking along chromosomes with super-resolution imaging, contact maps, and integrative modeling. bioRxiv. 2018; 374058. doi:10.1101/374058

98. Rand MD. A method of permeabilization of Drosophila embryos for assays of small molecule activity. J Vis Exp JoVE. 2014. doi:10.3791/51634

99. Rand MD, Kearney AL, Dao J, Clason T. Permeabilization of Drosophila embryos for introduction of small molecules. Insect Biochem Mol Biol. 2010;40: 792–804. doi:10.1016/j.ibmb.2010.07.007

100. . Erceg J, Abed JA, Goloborodko A, Lajoie BR, Fudenberg G, Abdennur N, et al. The genome-wide, multi-layered architecture of chromosome pairing in early Drosophila embryos. bioRxiv. 2018; 443028. doi:10.1101/443028

101. Schindelin J, Arganda-Carreras I, Frise E, Kaynig V, Longair M, Pietzsch T, et al. Fiji: an open-source platform for biological-image analysis. Nat Methods. 2012;9: 676–682. doi:10.1038/nmeth.2019

