## Supplementary_files.pdf for "Pericentromeric heterochromatin is hierarchically organized and spatially contacts H3K9me2 islands in euchromatin"

### Figure S1. Flow chart for identification of PCH Hi-C reads

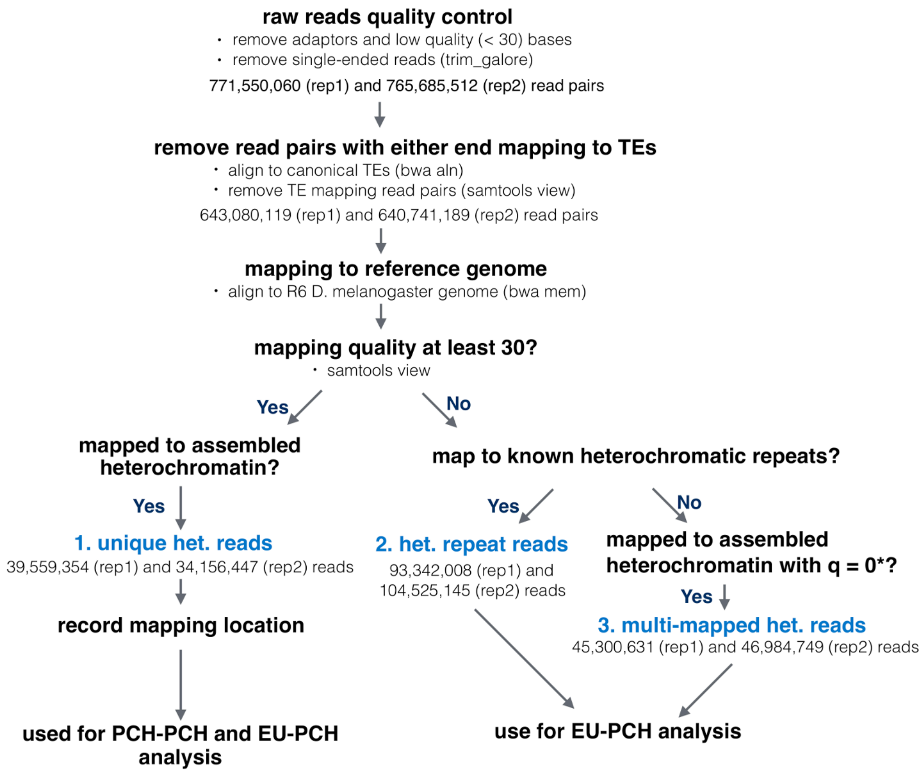

**Figure S2. Heatmap for the number of Hi-C read pairs supporting the spatial interactions between pairs of 100kb PCH windows for Hi-C replicate 2.** Note that only the PCH regions are shown.

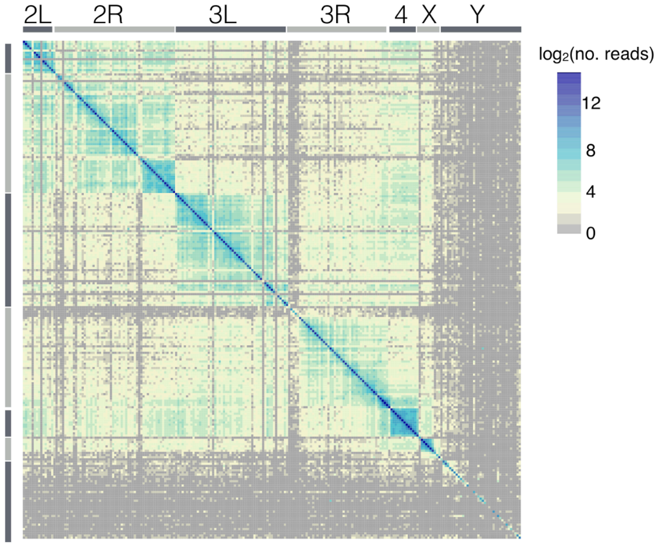

**Figure S3. Circular plot showing inter-arm and inter-chromosomal interactions.** Circular plot showing inter-arm and inter-chromosome interactions supported by 95, 99, and 99.9 percentile of Hi-C reads. Average mappability of each window is shown in the inner track.

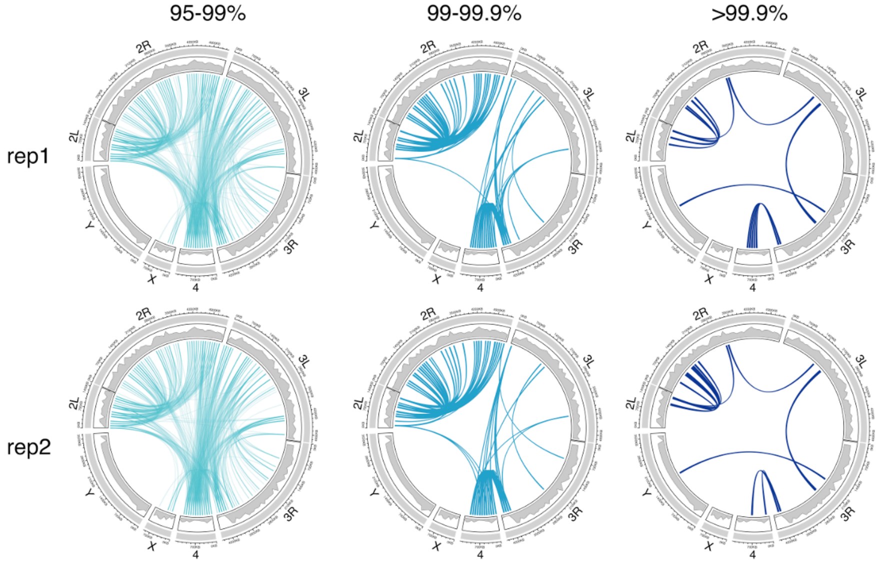

**Figure S4. Genome-wide normalized contact map of replicate 1.** Both unique euchromatic and PCH regions are shown. Blue bars are PCH regions while gray bars are euchromatic regions. Centromeres are denoted as triangles. Each element in the matrix represents the *log ratio* between the number of observed contact (Hi-C read pairs) and the number expected under the assumption that each 500kb window would have equal number of total interactions across the genome. The number of observed contacts involving Y chromosome is too low for proper normalization and thus excluded from representation in the figure. Note that this normalization may be biased against interactions involving PCH regions (EU-PCH and PCH-PCH) because much fewer reads uniquely mapped to PCH regions than euchromatic regions.

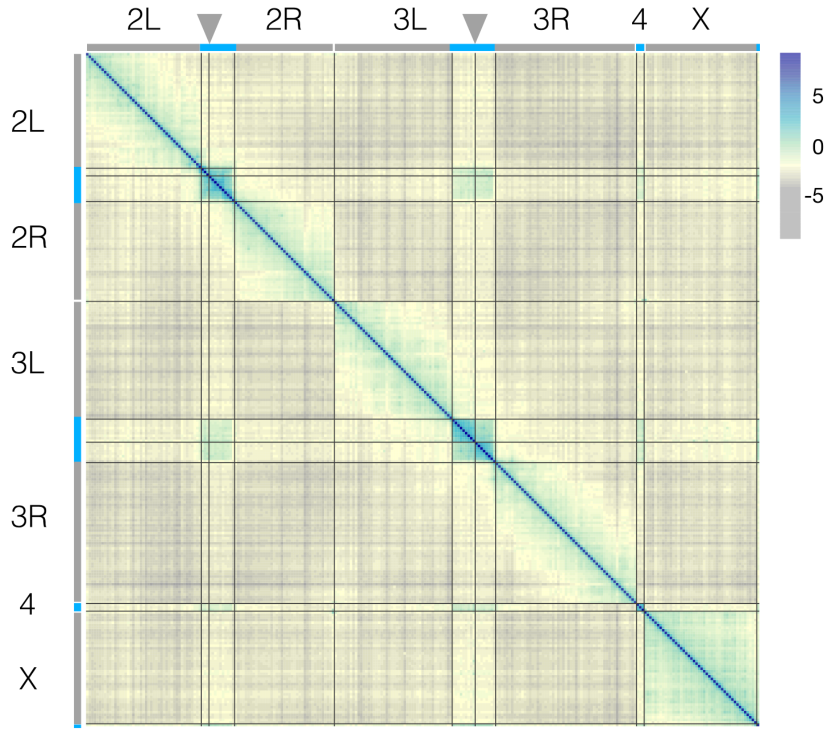

**Figure S5. Genome-wide normalized contact map of replicate 2.**

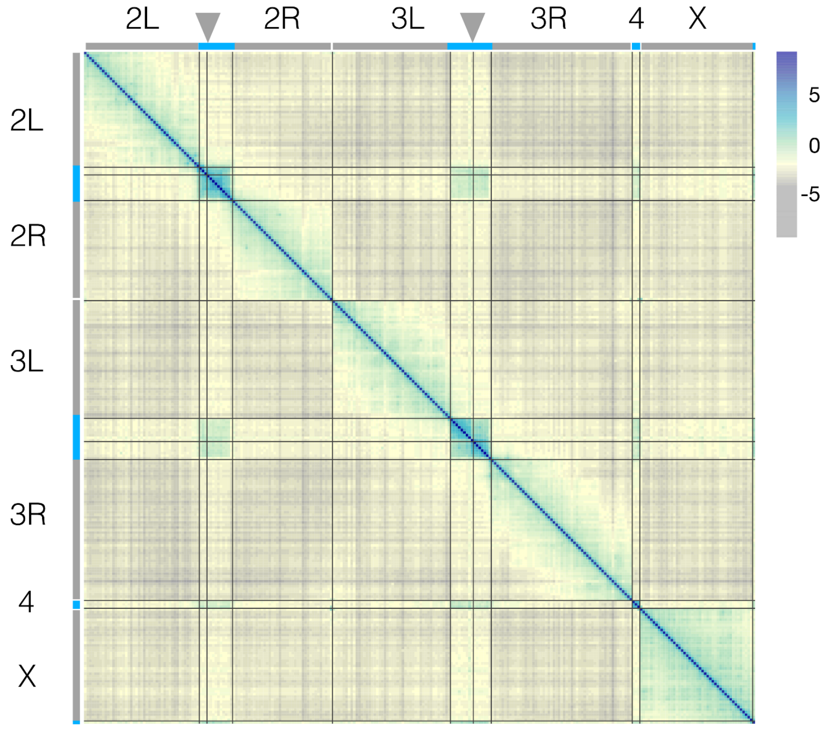

**Figure S6. FISH validation for identified PCH-PCH interaction using absolute distance.** (A) Boxplot and (B) histograms showing the absolute distance between PCH foci. Comparisons of distance between pairs of foci were tested with *Mann-Whitney test* (*p-values* in (A)) and *Fisher's exact test* (for proportion of overlapping foci, *p-values* = 0.001 (3R-4 vs 2R-4) and 0.22 (3R-4 vs 3R-2R)). Threshold for nuclei with overlapping foci is denoted with arrow, which corresponds to 0.1 μm.

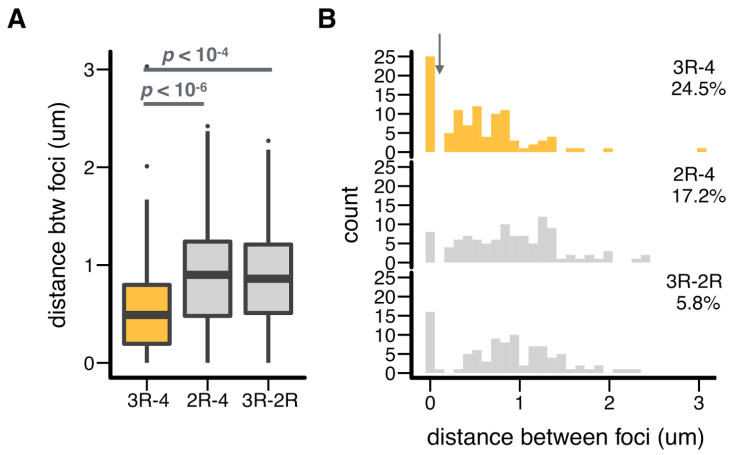

**Figure S7. Boxplot of the linear distance to PCH for H3K9me2 islands.** H3K9me2-enriched with and without PCH interactions are in green and gray respectively.

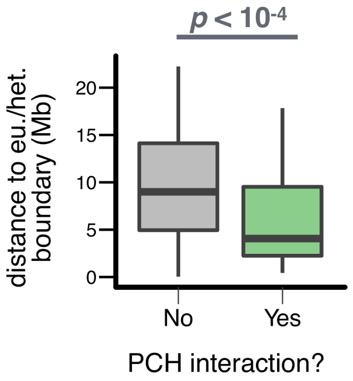

**Figure S8. Percentage of uniquely mapped heterochromatic Hi-C reads coming from a particular chromosome for euchromatic regions on different chromosomes.** Data for replicate 2 is shown.

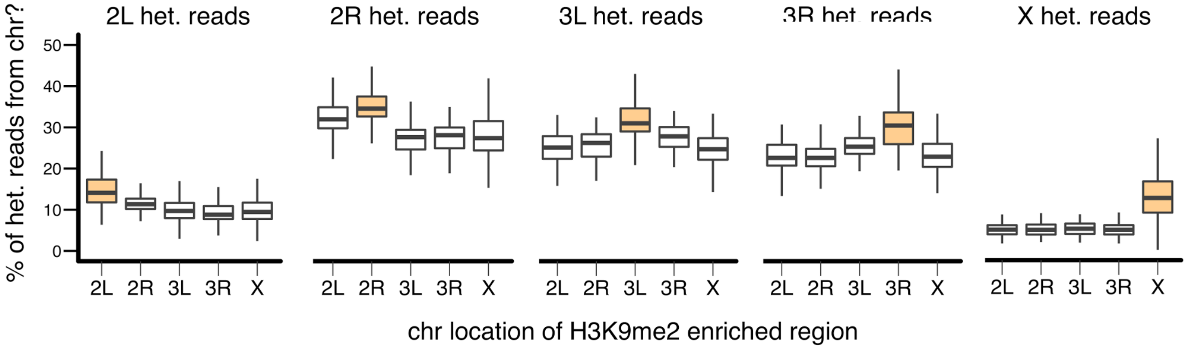

**Figure S9. H3K9me2 enrichment level for euchromatic regions chosen for FISH analysis.** There is H3K9me2 enrichment in both ORw1118 and wildtype strains for EU1-3, but none for control regions c.EU1-3. The fourth tracks (below RAL360, blue) are broad peaks called by Macs2 in ORw1118.

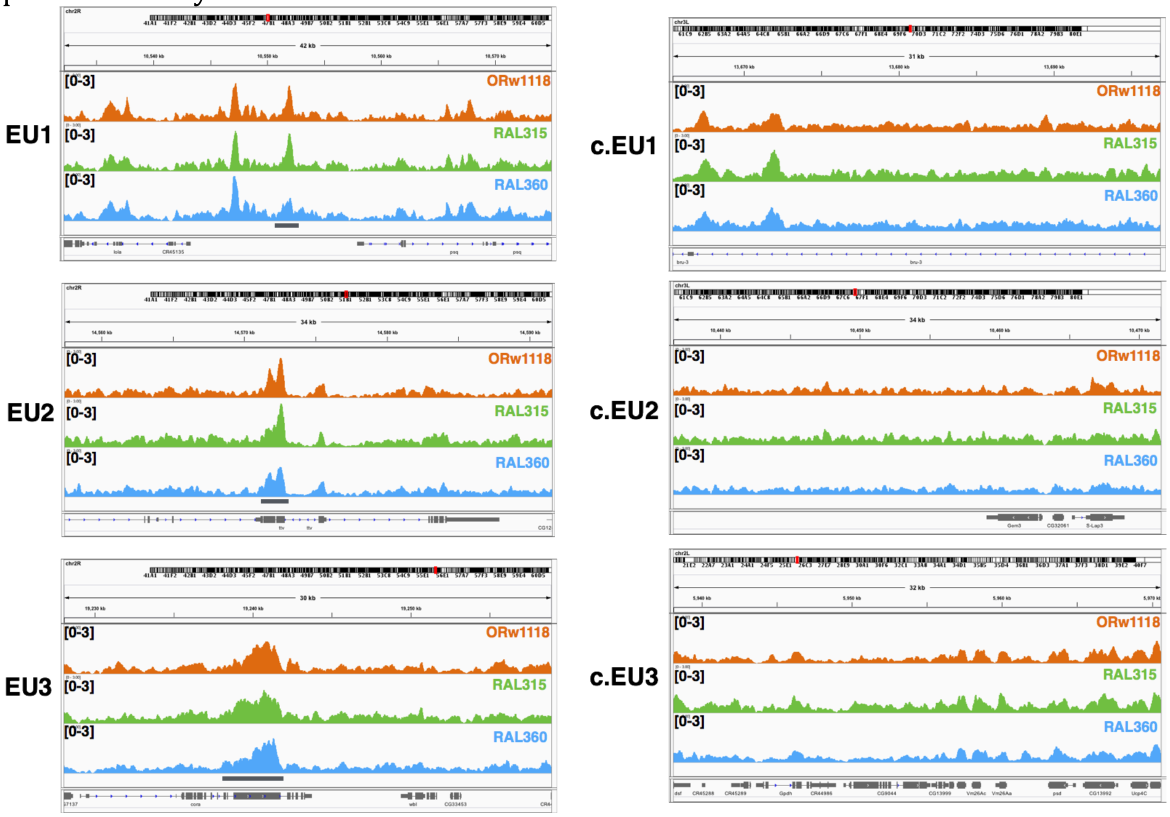

**Figure S10. Representative FISH images for chosen euchromatic regions and PCH.**

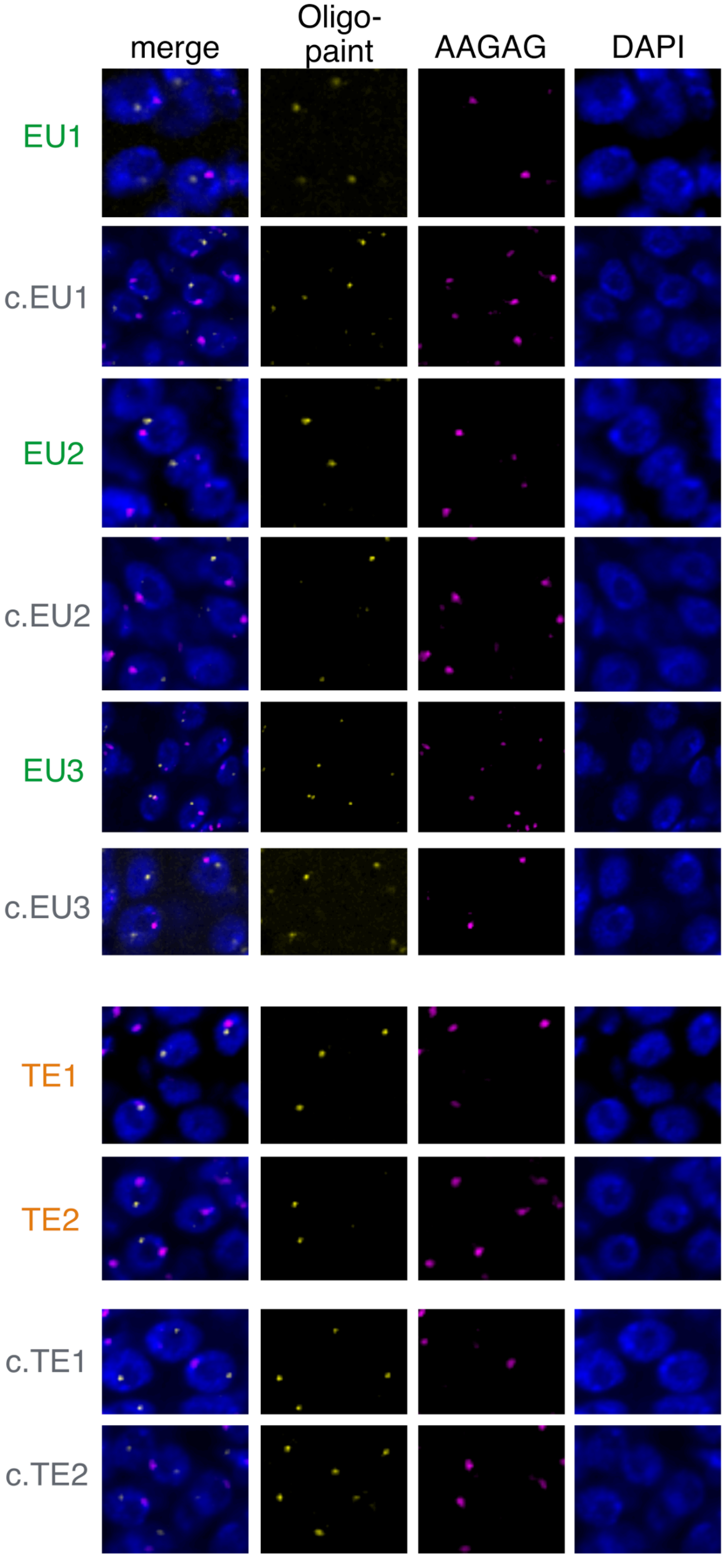

**Figure S11. FISH validation for identified EU-PCH interaction using absolute distance.** (A) Boxplot and (B) histogram showing the absolute distance between euchromatic loci and PCH. Comparisons of distance between pairs of foci were tested with *Mann-Whitney test* (*p-values* in (A)) and *Fisher's exact test* (for proportion of overlapping foci, *p-values* = 0.01 (EU1 vs c.EU1), 0.53 (EU2 vs c.EU2), and 0.15 (EU3 vs c.EU3)). Threshold for nuclei with overlapping foci is denoted with arrow, which corresponds to 0.1 µm.

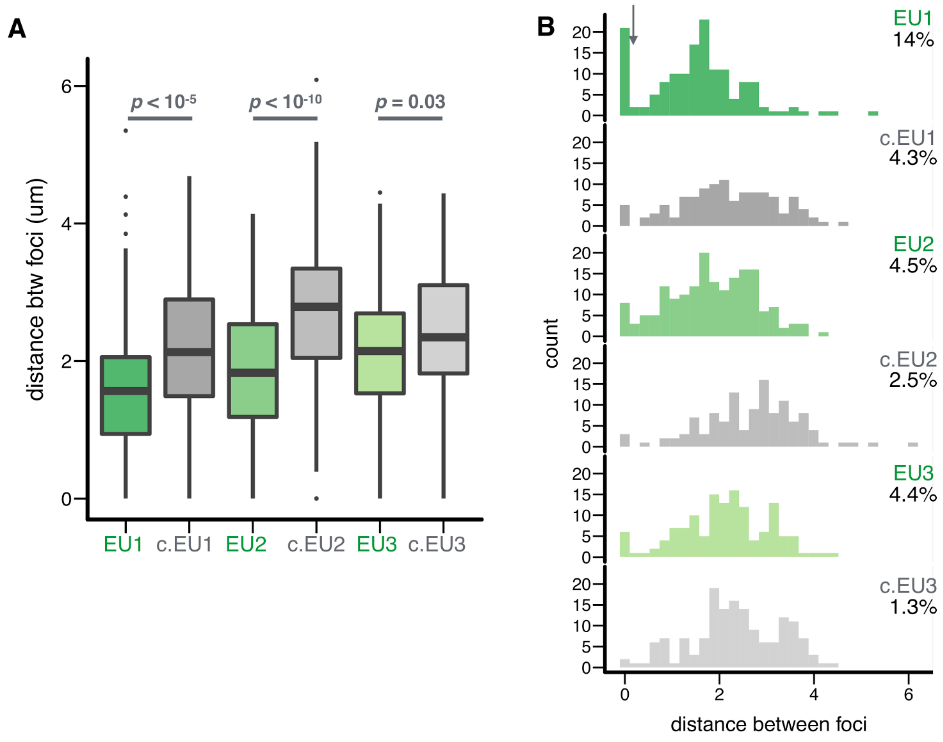

**Figure S12. H3K9me2 enrichment level in euchromatic TE neighborhood.** Strain-specific H3K9me2 enrichment was observed for TE1 and TE2. Third track (one below RAL315, green) shows the insertion position of TEs in ORw1118 predicted by TIDAL.

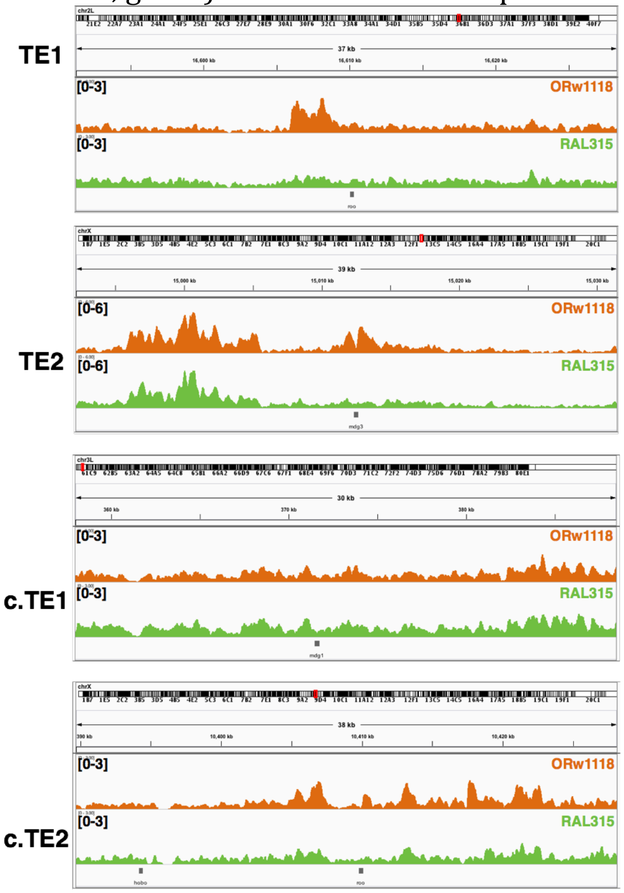

**Figure S13. Genomic distribution of TEs with and without PCH interactions.** The extent of local H3K9me2 enrichment at TEs is shown on the y-axis for TEs with (green) and without (gray) local H3K9me2 enrichment, and with (dark green) and without (light green) PCH interaction.

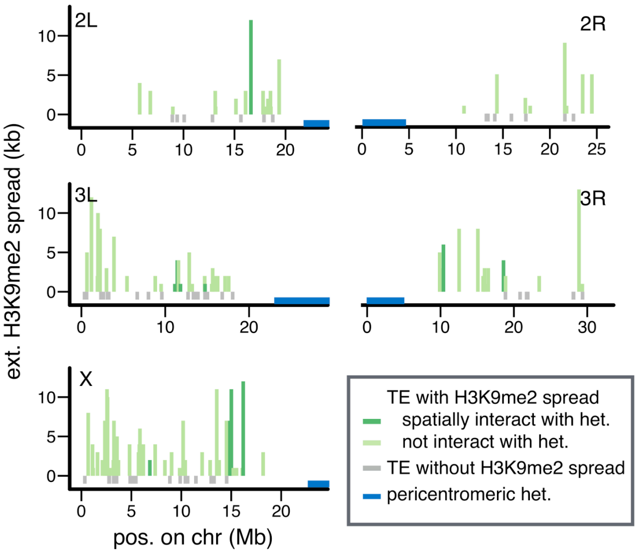

**Figure S14. Boxplot of the linear distance to PCH for euchromatic TEs.** TEs with and without PCH interactions are in dark and light green respectively.

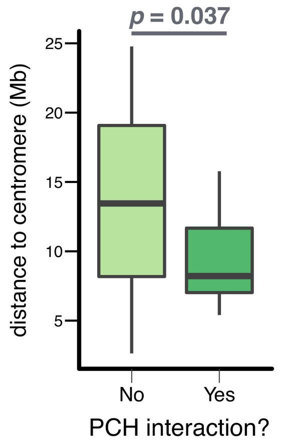

**Figure S15. FISH validation for identified TE-PCH interaction using absolute distance.** (A) Boxplot and (B) histogram showing the absolute distance between euchromatic TEs and PCH. Comparisons of distance between pairs of foci were tested with *Mann-Whitney test* (*p-values* in (A)) and *Fisher's exact test* (for proportion of overlapping foci, *p-values* = 0.0007 (TE1), 0.07 (TE2), 0.48 (c.TE1), and 1 (c.TE2)). Threshold for nuclei with overlapping foci is denoted with arrow, which corresponds to 0.1 µm.

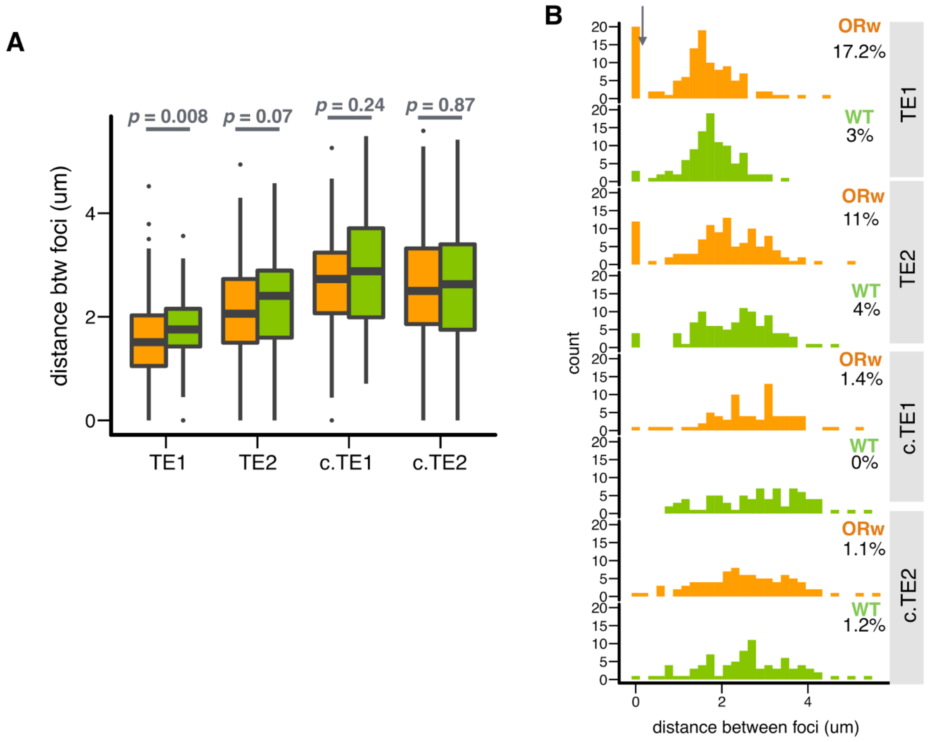

**Figure 16. Representative FISH images for permealized embryos (EPS) and permealized embryos with 1,6-hexanediol treatment (EPS+HD)**

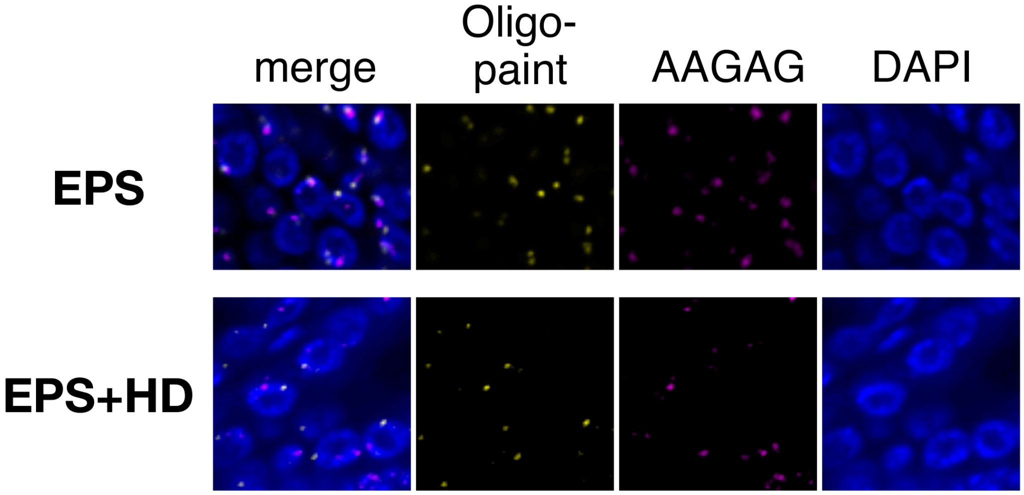

**Figure S17. FISH validation for the influence of 1,6-Hexanediol on the spatial associations between euchromatic TE and PCH using absolute distance.** (A) Boxplot and (B) histogram showing the absolute distance between euchromatic TE and PCH. Comparisons of distance between pairs of foci were tested with *Mann-Whitney test* (*p-values* in (A)) and *Fisher's exact test* (for proportion of overlapping foci, *p-values* = 0.02 (ORw, EPS vs EPS+HD), 1 (WT, EPS vs EPS+HD), 0.057 (ESP treatment, ORw vs WT), 0.55 (HD treatment, ORw vs WT)). Threshold for nuclei with overlapping foci is denoted with arrow, which corresponds to 0.1 μm.

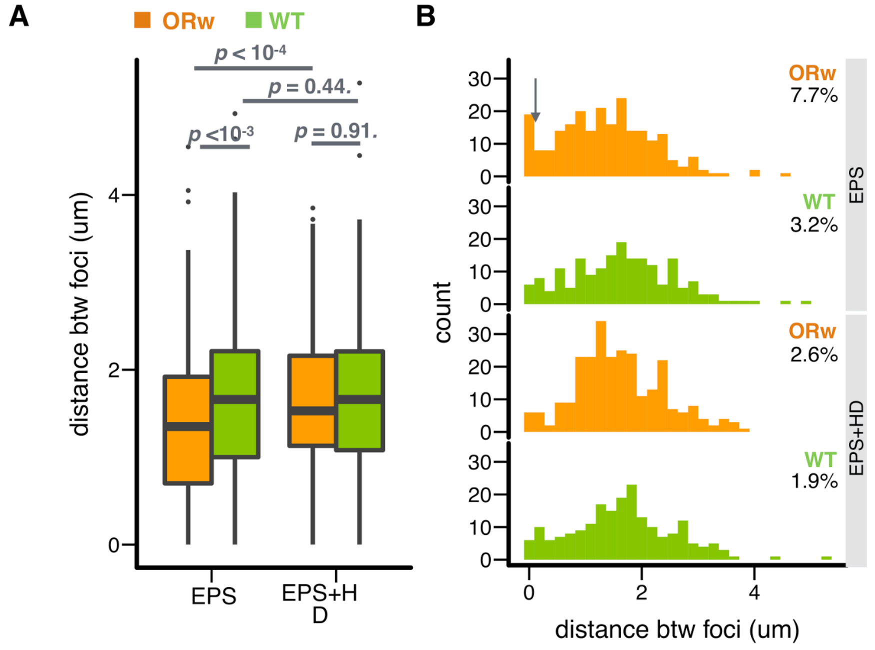

**Figure S18. Extent and magnitude of H3K9me2 enrichment of TEs with and without PCH interaction.**

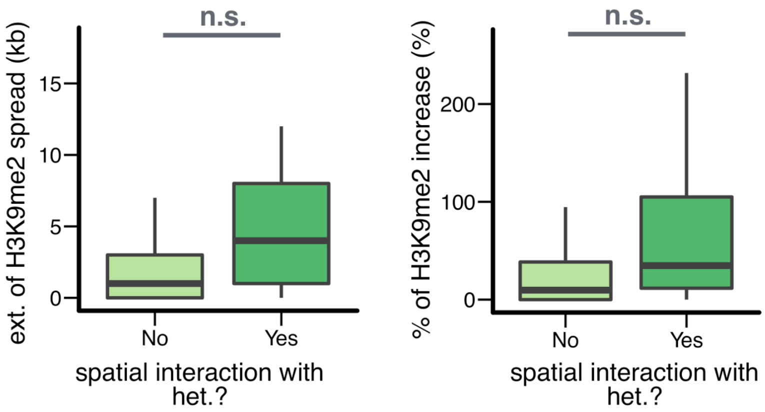

**Figure S19. IDR plots for ORw1118**

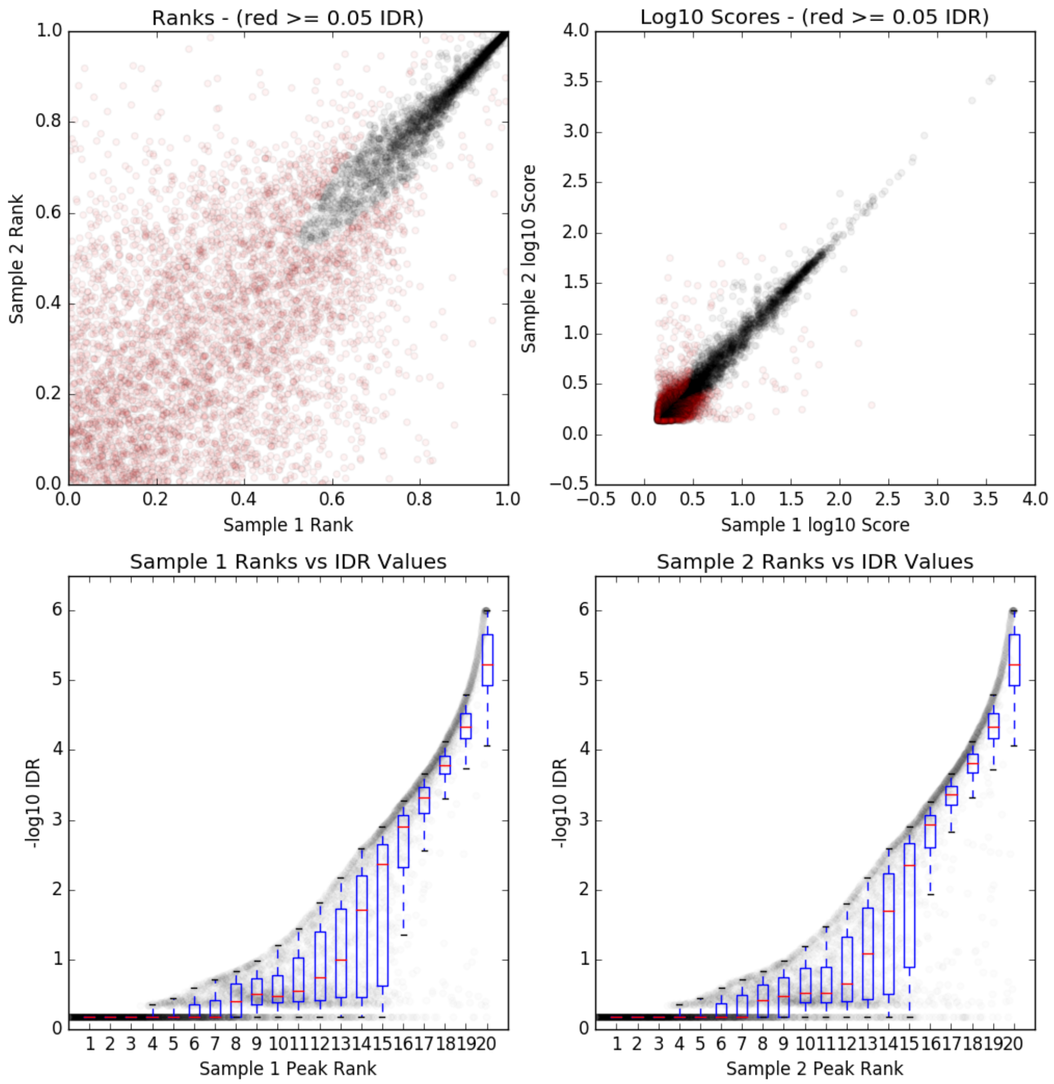

**Figure S20. IDR plots for RAL315**

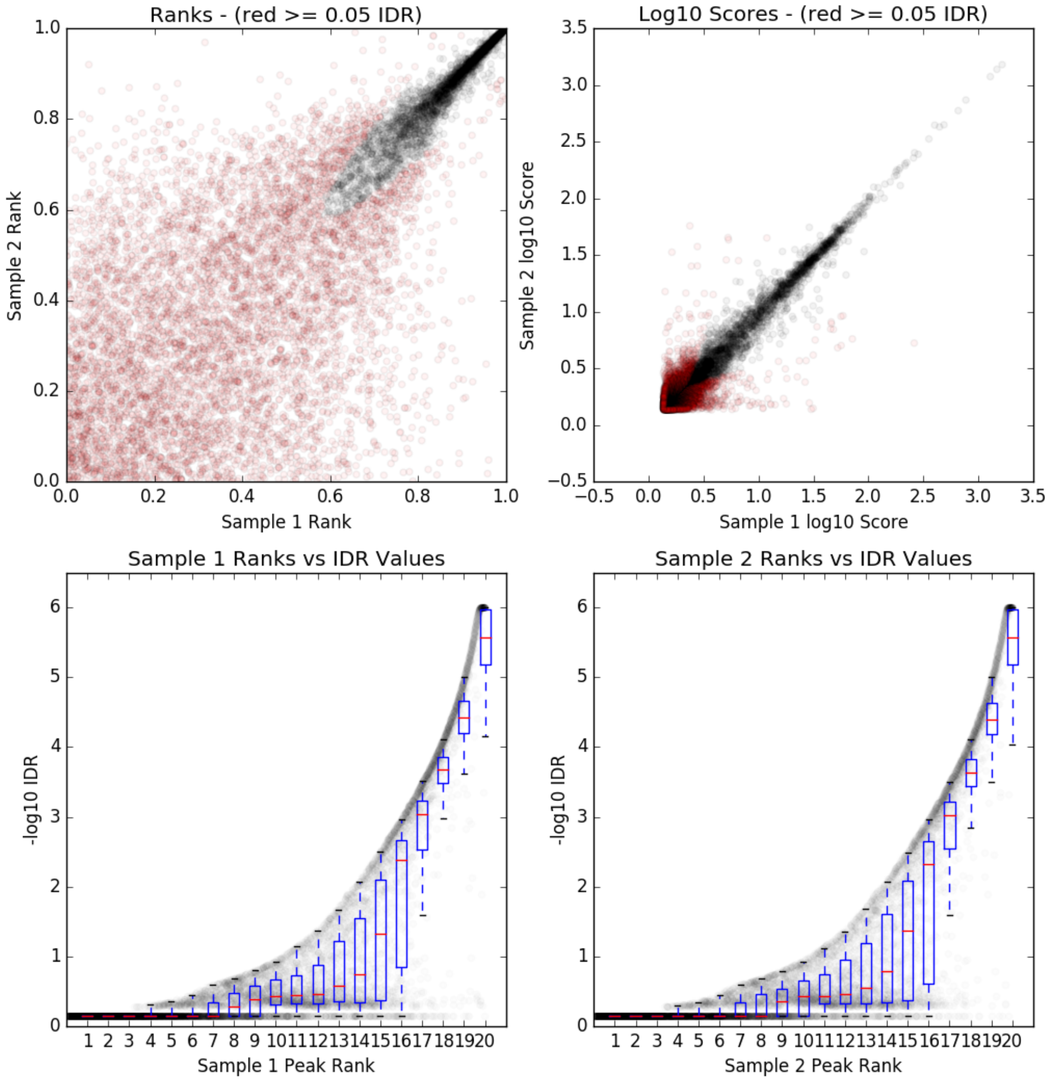

**Figure S21. IDR plots for RAL360**

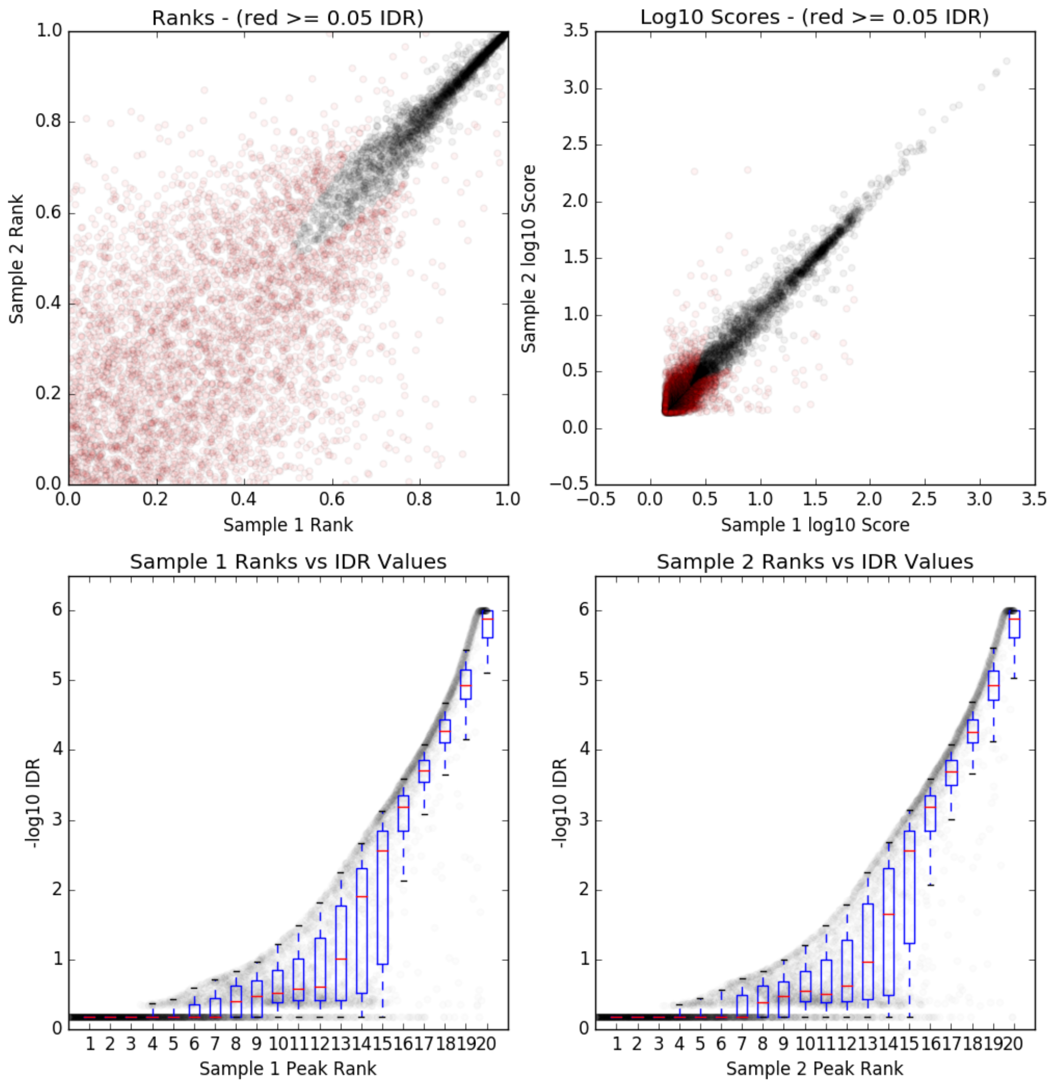

**Figure 22. X-Y plots for the estimated proportion of euchromatin-PCH reads, and the associated *p-values*, between Hi-C replicates for H3K9me2-enriched regions.**

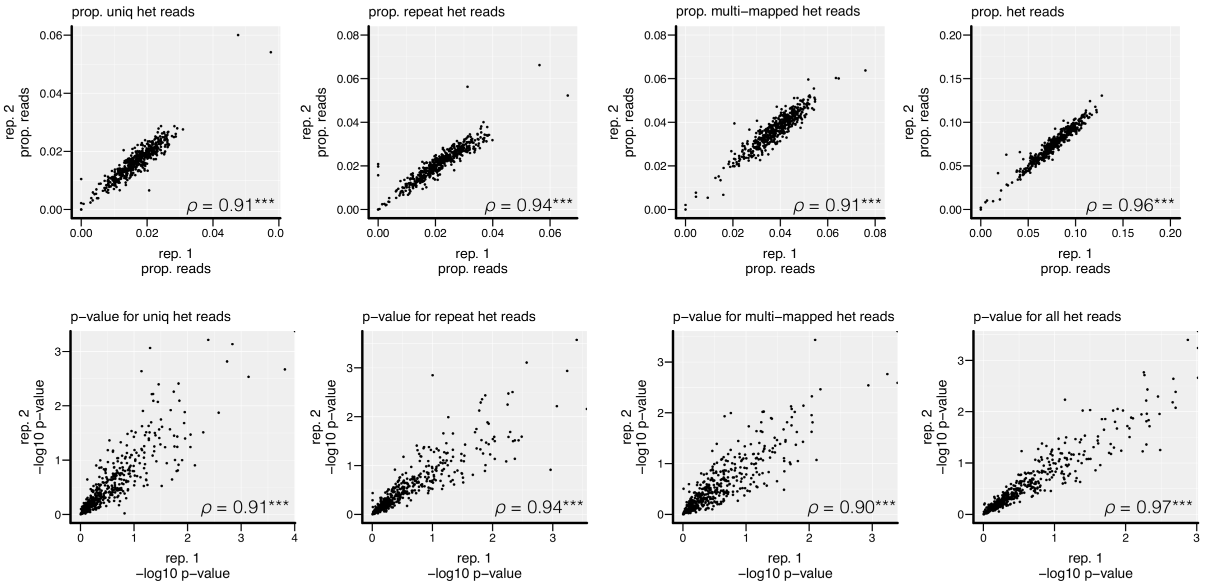

**Figure S23. X-Y plots for the estimated proportion of euchromatin-PCH reads, and the associated *p-values*, between Hi-C replicates for TEs.**

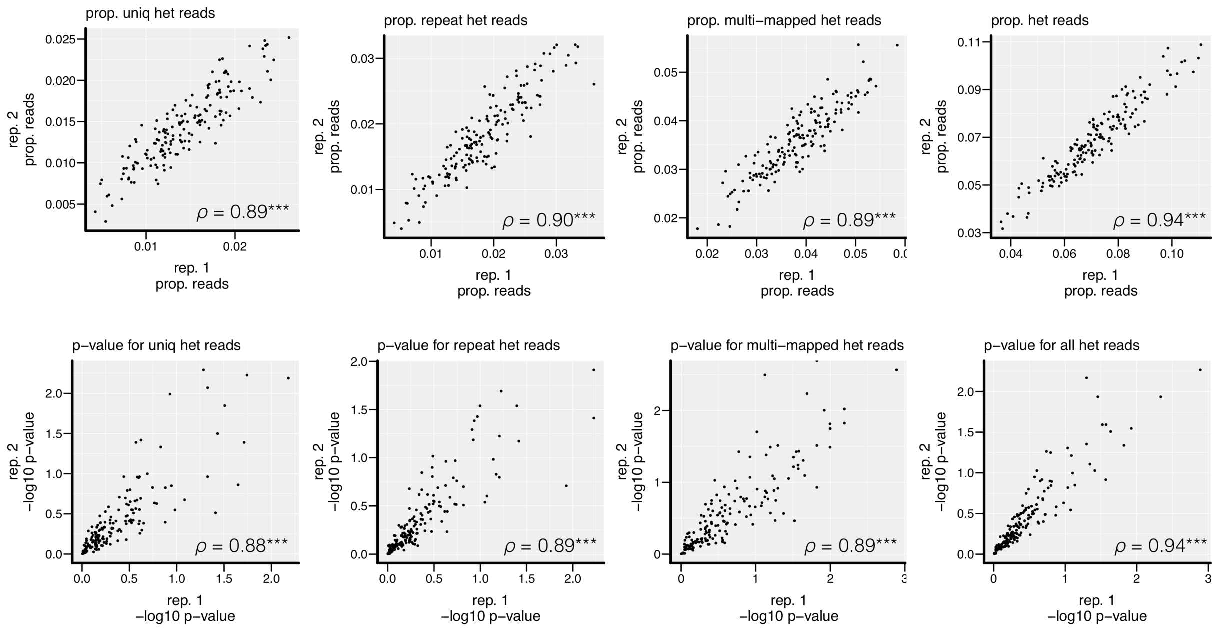

**Figure S24. Distribution of estimated euchromatin-PCH read pairs for random regions on different chromosomes.**

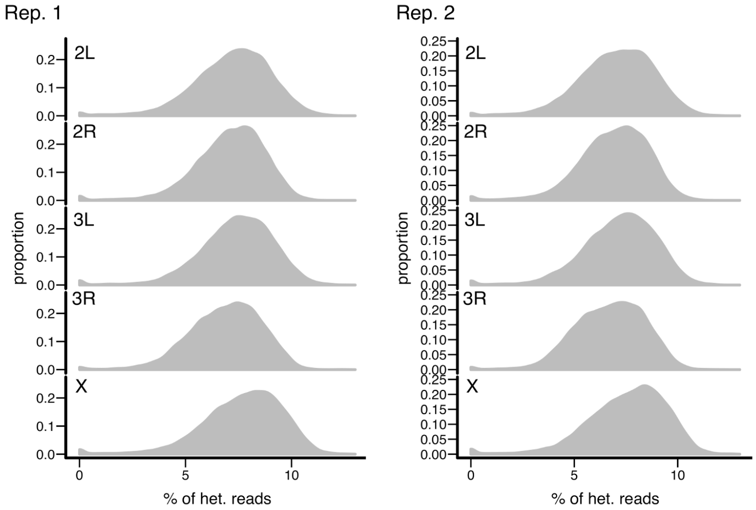

**Figure S25. Distribution of estimated euchromatin-PCH read pairs for random regions of different size.**

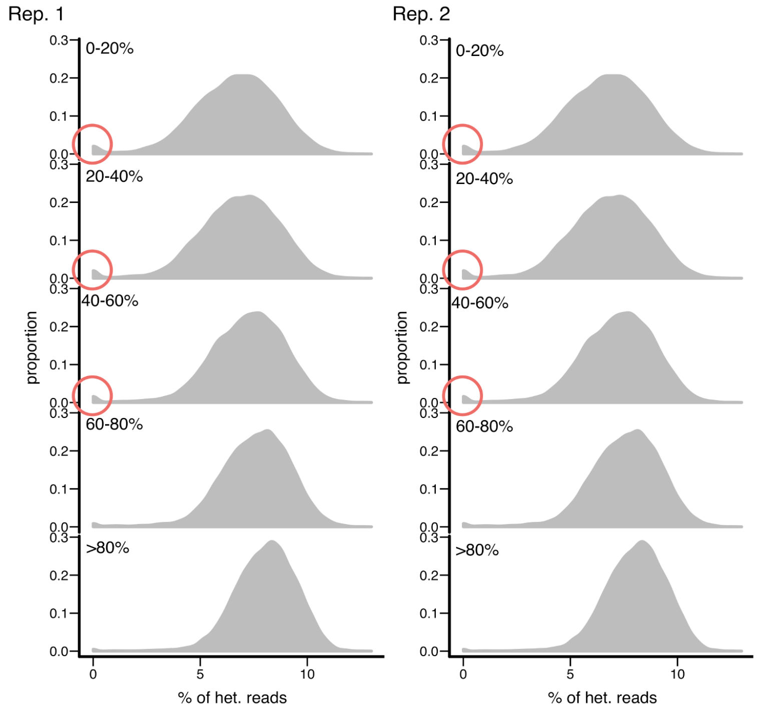

**Table S1. List of heterochromatic simple repeats**

| **Repeat** | **Note** |
|---|---|
| 1.686 | |
| 1.688 | |
| AAAACAT | |
| AAAAG | |
| AAAATAT | AATAT variant |
| AAACAAT | |
| AAACAC | AACAC variant |
| AAAGAC | AAGAC variant |
| AAATTACT | |
| AACAC | 1.672 |
| AAGAC | 1.689, 1.701 |
| AAGACATGAC | |
| AAGACTAGAC | |
| AAGAG | 1.705 |
| AAGAGAAGAGAG | AAGAG variant |
| AAGAGAG | AAGAG variant |
| AAGAGG | AAGAG variant |
| AAGGAG | AAGAG variant |
| AATAC | 1.68 |
| AATAG | 1.693 |
| AATAGAC | 1.688 |
| AATAT | 1.672 |
| AATATAT | AATAT variant |
| ACATATAT | |
| ACCAGTACGGG | undeca |
| ACCGAGTACGGG | dodeca |
| AGATG | |
| His1 | |
| His2A | |
| His2B | |
| His3 | |
| His4 | |
| rDNA | |
| Rsp | |

**Table S2. Number of Hi-C read pairs for pairs of PCH regions**

| | | no. of read pairs | | % of all read pairs | |
|---|---|---|---|---|---|
| chr 1 | chr 2 | replicate 1 | replicate 2 | replicate 1 | replicate 2 |
| 2L | 2L | 1164391 | 773042 | 9.66% | 9.63% |
| 2L | 2R | 37392 | 38783 | 0.31% | 0.48% |
| 2L | 3L | 9525 | 9277 | 0.08% | 0.12% |
| 2L | 3R | 7190 | 6802 | 0.06% | 0.08% |
| 2L | 4 | 6822 | 6949 | 0.06% | 0.09% |
| 2L | X | 1385 | 1408 | 0.01% | 0.02% |
| 2L | Y | 421 | 452 | 0.00% | 0.01% |
| 2R | 2R | 2758050 | 1778783 | 22.87% | 22.15% |
| 2R | 3L | 26163 | 24688 | 0.22% | 0.31% |
| 2R | 3R | 20945 | 20818 | 0.17% | 0.26% |
| 2R | 4 | 19009 | 18245 | 0.16% | 0.23% |
| 2R | X | 4303 | 4177 | 0.04% | 0.05% |
| 2R | Y | 1491 | 1472 | 0.01% | 0.02% |
| 3L | 3L | 2490763 | 1600901 | 20.66% | 19.94% |
| 3L | 3R | 42885 | 43355 | 0.36% | 0.54% |
| 3L | 4 | 22377 | 21738 | 0.19% | 0.27% |
| 3L | X | 3526 | 3532 | 0.03% | 0.04% |
| 3L | Y | 1432 | 1416 | 0.01% | 0.02% |
| 3R | 3R | 2492544 | 1731036 | 20.67% | 21.56% |
| 3R | 4 | 17962 | 17364 | 0.15% | 0.22% |
| 3R | X | 3348 | 3251 | 0.03% | 0.04% |
| 3R | Y | 1649 | 1765 | 0.01% | 0.02% |
| 4 | 4 | 2050167 | 1321436 | 17.00% | 16.46% |
| 4 | X | 2414 | 2552 | 0.02% | 0.03% |
| 4 | Y | 771 | 678 | 0.01% | 0.01% |
| X | X | 778807 | 537628 | 6.46% | 6.69% |
| X | Y | 229 | 231 | 0.00% | 0.00% |
| Y | Y | 91377 | 58732 | 0.76% | 0.73% |

**Table S3. Chromatin environment of euchromatic H3K9me2-enriched regions interacting with PCH**

| Sexton et al. 2012 | | | | | Active vs other | | Null vs other | |
|---|---|---|---|---|---|---|---|---|
| 3D interaction with PCH | Active | HP1a-enriched | Null | Polycomb-enriched | FET *p-value* | odds ratio | FET *p-value* | odds ratio |
| Yes | 18 | 1 | 20 | 4 | 7.76E-03 | 2.51 | 3.05E-02 | 0.48 |
| No | 100 | 3 | 291 | 57 | | | | |

| Filon et al. 2010 | | | | | | Red+Yellow vs other | | Black vs other | |
|---|---|---|---|---|---|---|---|---|---|
| 3D interaction with PCH | BLACK | BLUE | GREEN | RED | YELLOW | FET *p-value* | odds ratio | FET *p-value* | odds ratio |
| Yes | 12 | 12 | 2 | 0 | 16 | 2.10E-02 | 2.22 | 5.19E-04 | 0.30 |
| No | 260 | 82 | 13 | 17 | 81 | | | | |

| Kharchenko et al. 2011 | | | | | | | | | | 1,2,3,4 vs other | | 9 vs other | |
|---|---|---|---|---|---|---|---|---|---|---|---|---|---|
| 3D interaction with PCH | 1 | 2 | 3 | 4 | 5 | 6 | 7 | 8 | 9 | FET *p-value* | odds ratio | FET *p-value* | odds ratio |
| **S2 cells** | | | | | | | | | | | | | |
| Yes | 4 | 5 | 1 | 3 | 5 | 1 | 0 | 0 | 5 | 5.27E-05 | 6.14 | 8.64E-03 | 0.26 |
| No | 9 | 19 | 3 | 19 | 34 | 8 | 3 | 11 | 106 | | | | |
| **BG3 cells** | | | | | | | | | | | | | |
| Yes | 2 | 4 | 2 | 3 | 4 | 0 | 0 | 5 | 5 | 1.13E-02 | 3.05 | 7.85E-01 | 1.16 |
| No | 10 | 9 | 13 | 21 | 22 | 11 | 3 | 125 | 46 | | | | |

**Table S4. Properties of euchromatic H3K9me2-enriched regions interacting with PCH**

| | 3D interaction criteria | | | |
|---|---|---|---|---|
| | **both rep. sig.** *p-values* | **either rep.sig** *p-values* | **test type** | **Note** |
| Distance to centromere | **5.0E-02** | **2.0E-04** | *Mann-Whitney U* | with 3D interaction shorter |
| proportion of coding sequence | **4.7E-03** | **7.6E-04** | *Mann-Whitney U* | with 3D interaction higher |
| X vs autosome | 3.1E-01 | 2.7E-01 | *Fisher's Exact* | |
| size of H3K9me2 enriched region | 3.4E-01 | 3.1E-01 | *Mann-Whitney U* | |
| H3K9me2 enrichment | 9.5E-01 | 9.1E-01 | *Mann-Whitney U* | |
| enrichment of Active TAD (Sexton et al. 2012) | **7.8E-03** | **3.2E-02** | *Fisher's Exact* | odds ratio = 1.88 (either) |
| enrichment of Red and Yellow chromatin (Filion et al. 2010) | **2.1E-02** | 1.2E-01 | *Fisher's Exact* | odds ratio = 1.59 (either) |
| enrichment of 1-4 of modEncode 9 states, S2 (Kharchenko et al. 2011) | **5.3E-05** | **1.6E-02** | *Fisher's Exact* | odds ratio = 2.55 (either) |
| enrichment of 1-4 of modEncode 9 states, BG3 (Kharchenko et al. 2011) | **1.1E-02** | **3.8E-02** | *Fisher's Exact* | odds ratio = 2.20 (either) |
| depletion of Null TAD (Sexton et al. 2012) | **3.1E-02** | **1.0E-02** | *Fisher's Exact* | odds ratio = 0.50 (either) |
| depletion of Black chromatin (Filion et al. 2010) | **5.2E-04** | **6.0E-04** | *Fisher's Exact* | odds ratio = 0.40 (either) |
| depletion of 9 of modEncode 9 states, S2 (Kharchenko et al. 2011) | **8.6E-03** | **7.3E-03** | *Fisher's Exact* | odds ratio = 0.35 (either) |
| depletion of 9 of modEncode 9 states, BG3 (Kharchenko et al. 2011) | 7.9E-01 | 3.8E-01 | *Fisher's Exact* | |

**Table S5. Oligopaint targeted regions**

| ID | chr | window start | window end | no. probes | probe density (per kb) | distance to centromere (Mb) | Notes |
|---|---|---|---|---|---|---|---|
| TE1 | 2L | 16591247 | 16628353 | 504 | 13.58 | 2.96 | roo |
| TE2 | X | 14992136 | 15031508 | 505 | 12.83 | 3.92 | mdg3 |
| c.TE1 | 3L | 357967 | 388478 | 507 | 16.62 | 18.07 | mdg1 |
| c.TE2 | X | 10389680 | 10428155 | 507 | 13.18 | 8.52 | roo |
| EU1 | 2R | 10532124 | 10575010 | 503 | 11.73 | 1.69 | |
| EU2 | 2R | 14557395 | 14591631 | 503 | 14.69 | 5.71 | |
| EU3 | 2R | 19228022 | 19258953 | 507 | 16.39 | 10.38 | |
| c.EU1 | 3L | 13665384 | 13696922 | 507 | 16.08 | 4.73 | |
| c.EU2 | 3L | 10436331 | 10471601 | 505 | 14.32 | 7.98 | |
| c.EU3 | 2L | 5938111 | 5970628 | 508 | 15.62 | 13.62 | |

**Table S6. Properties of euchromatic TEs interacting with PCH**

| | **3D interaction criteria** | | | |
|---|---|---|---|---|
| | **both rep. sig.** *p-values* | **either rep.sig** *p-values* | **test type** | **Note** |
| Distance to centromere | **2.69E-02** | **3.73E-02** | *Mann-Whitney U* | with 3D interaction shorter |
| X vs autosome | 1.00E+00 | 1.00E+00 | *Fisher's Exact* | |
| extent of H3K9me2 spread | 1.47E-01 | 3.03E-01 | *Mann-Whitney U* | |
| % increase in H3K9me2 enrichment | 1.90E-01 | 5.34E-01 | *Mann-Whitney U* | |
| type of TEs (TIR, non-LTR, LTR) | 5.06E-01 | 9.84E-02 | *Chi-square* | |
| DNA vs RNA TEs | 6.33E-01 | 2.39E-01 | *Fisher's Exact* | |
| Population frequencies | **1.03E-02** | **4.19E-03** | *Student t test* | with 3D interaction lower |

#### Supplementary Text

##### *Trans* epigenetic effects of TEs

We proposed that the spatial interactions between euchromatic loci and PCH on one chromosome could influence the chromatin environment of the homolog, such as *trans*-silencing, due to strong somatic pairing in *Drosophila* (see main text). To address this question, we focused on polymorphic TE insertions between genomes and investigate whether they could lead to *trans*-epigenetic effects, or influencing the chromatin of the homolog, when in heterozygous states. We analyzed the enrichment of H3K9me2 around heterozygous TE insertions in two wildtype strains and their F1 offspring (from both directions of the cross). Maternal and paternal alleles were distinguished using previously identified SNPs in the two strains *(80)*. We modified the approach in *(34)*, which compares the H3K9me2 enrichment level of *all sites* in a window between strains with and without TE insertions. Instead, we only compared the H3K9me2 enrichment level of *SNPs,* and the *trans* effects would be confirmed if the H3K9me2 fold enrichment levels for both alleles at *SNPs* in the F1 (with and without TEs) are higher than in the parental strain without the TE.

Eight and 17 TEs have significant signals of *trans*-epigenetic effects in either direction of the crosses, which accounted for 7.9% and 16.8% of all TEs analyzed. **Supplementary Text -Figure 1 (left)** shows an exemplar TE with *trans*-epigenetic effects. While H3K9me2 enrichment is only found in parental strain with TE insertion, such enrichment was found for *both* alleles, with and without TE insertion, in the F1 offspring. In contrast, **Supplementary Text - Figure 1 (right)** shows an exemplar TE without *trans*-epigenetic effects. Intriguingly, we did not find TEs with significant *trans*-epigenetic effects for both direction of the cross. However, given the high false negative rates of analysis based on SNP alone (see below), it is still inconclusive if there are strong maternal effects for the observed *trans*-epigenetic effects. Unlike TEs identified as spatially interacting with PCH (see main text), we did not find differences in the distance to PCH between TEs with and without *trans*-epigenetic effects. Many factors could have contributed to this. For example, the strength of homolog pairing varies across the genome *(71)*, which could also influence the tendency of a TE to show *trans*-epigenetic effects.

It is worth noting there is on average ~20 SNPs within 1kb window, which suggests lower statistical power for analysis based on *SNPs* alone than that based on *all sites*. SNPs that are within 100bp could highly likely be covered by the same sequencing read and thus contain redundant information for inference. Randomly sampling SNPs that are within 100bp led to even fewer SNPs with unique information in a 1kb window (~5). Indeed, we found a high false negative rate (46.29%) by comparing results based on *SNP-based* and *all-sites* approach for identifying TE-induced H3K9me2 enrichment in the *parental strain*. Future analysis involving more strains that have larger genetic divergence, coupled with Hi-C studies and homolog pairing maps, will help further address the potential functional consequence of euchromatin-PCH 3D contacts.

**Supplementary Text - Figure 1. H3K9me2 enrichment level around exemplar TEs in parental strains (P) and F1.** A TE on 3R (left) was found to have *trans*-epigenetic effects while another TE on the 3L (right) was not. Window size 2kb, with LOESS smoothing (alpha = 0.05).

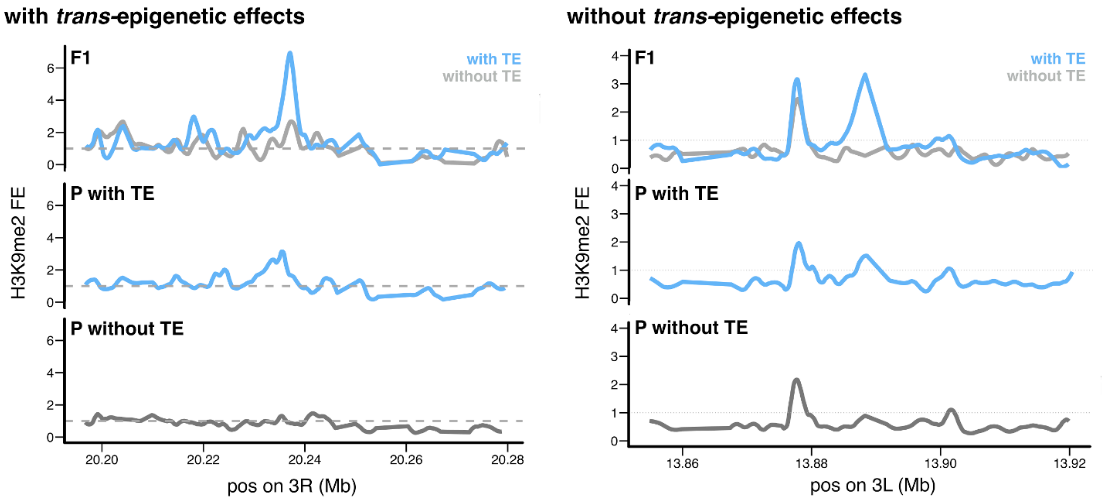


**Supplementary Text - Methods**

We performed ChIP-seq targeting H3K9me2 using 16-18hr embryos of RAL315, RAL360, and the F1 of the two wildtype strains. TE insertion positions in these two strains were from *(80)* and we only included TEs with coverage ratio at least three, which is a highly stringent threshold. The conversions between release 5 coordinates (SNPs, see below) and release 6 coordinates (TE insertion positions) were done with liftOver (https://genome.ucsc.edu).

We downloaded genomes of RAL315 and RAL360 from Drosophila Genome Nexus (in release 5, *(89)*) and followed the recommendations by masking following regions that are error-prone: 3bp around indels, regions of identity by descent, and regions with residual heterozygosity. After such filtering, we generated a list of SNP that have different allelic states between RAL315 and RAL360. Raw reads from ChIP-seq experiments were pre-processed as methods described in main text. Filtered reads were mapped to *combined genomes of RAL315 and RAL360* using bwa mem with default parameters. We generated pileup tables, which contain counts for alternative alleles, using samtools' mpileup command with mapping quality cutoff 20. We filtered out SNPs whose alternative allele (e.g. RAL360 allele in RAL315) has more than one supporting reads *in the parental strains* (i.e. potentially heterozygous SNPs in parental strains). For F1s, we only included SNPs whose combined read depth of two replicates were above three.

To estimate allelic-specific fold enrichment of H3K9me2 for each SNP, we first (1) normalized sequencing depth between IP and INPUT samples using total read depth and (2) divided the normalized read count of IP by normalized read count of INPUT sample. Because the background H3K9me2 enrichment levels are different between genotypes, we used median fold enrichment of *SNPs* in +/-20-40kb to normalize the background enrichment levels between genotypes. We used one-side *Mann-Whitney U test* to investigate whether the fold enrichment of *SNPs* in (1) the parental strain with TE, (2) with TE allele in F1 or (3) without TE allele in F1 is higher than that in the parental strain without TE (**Figure 2**). A TE is deemed as having *trans*-epigenetic effects if both F1 alleles with and without TEs have significantly higher H3K9me2 fold enrichment than that of the parental strain without TE. The window size used was defined by finding the extent of H3K9me2 spread from TEs using *all-sites method* as previously described (**Supplementary Text - Figure 2,***(34)*). TEs that do not have H3K9me2 enrichment in the parental strain with TE or have fewer than 10 surrounding SNPs were excluded from the analysis, and in total101 TEs were analyzed. All analysis combined RAL315-speicifc and RAL360-specific TEs together.

**Supplementary Text - Figure 2. *SNP-based* and *all-sites* based method for identifying the epigenetic effects of TEs.** Smoothed lines represent H3K9me2 fold enrichment across the genome (all sites), while vertical lines represent the H3K9me2 fold enrichment for SNPs.

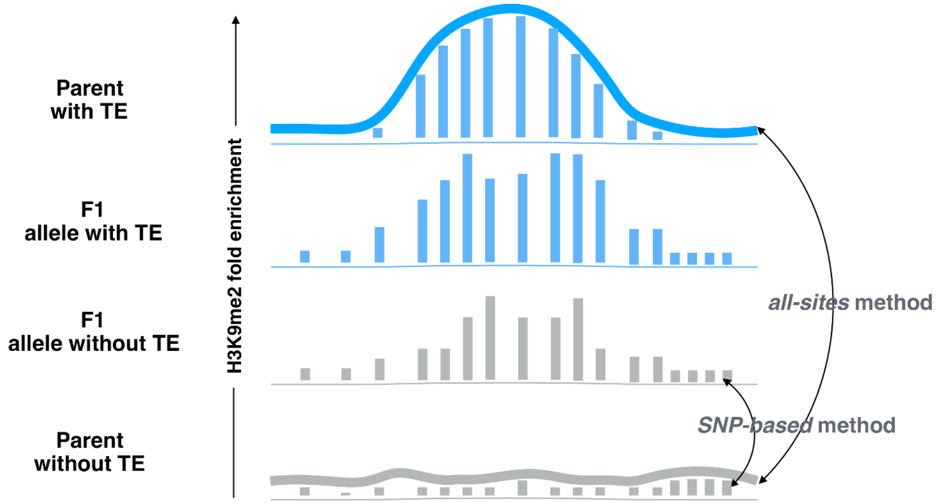
